# Uncovering the function of Wisp1 in whole-body glucose homeostasis: insights from Wisp1 knockout mice

**DOI:** 10.64898/2026.01.15.699691

**Authors:** Rebeca Fernandez-Ruiz, Ainhoa García-Alamán, Marta Fontcuberta-PiSunyer, Marta Perea-Atienzar, Jessica Angulo-Capel, Colleen K Hadley, Zeran Lin, Paul Cohen, Josep Vidal, Rosa Gasa

**Affiliations:** Institut d’Investigacions Biomèdiques August Pi i Sunyer (IDIBAPS), Barcelona, Spain; CIBER de Diabetes y Enfermedades Metabólicas Asociadas, Instituto de Salud Carlos III, Madrid, Spain; University of Barcelona, Barcelona, Spain; Laboratory of Molecular Metabolism, The Rockefeller University, New York, US; Weill Cornell/Rockefeller/Sloan Kettering Tri-institutional MD-PhD Program, New York, US; Endocrinology and Nutrition Department, Hospital Clinic of Barcelona, Barcelona, Spain

## Abstract

WNT1-inducible signaling pathway protein 1 (Wisp1/CCN4) is a matricellular protein implicated in inflammation and metabolic dysfunction in obesity, yet its role in whole-body glucose metabolism remains unclear. In this study, Wisp1 knockout (KO) mice were analysed under physiological and high-fat (HF) diet conditions to define its impact on metabolic regulation. Neither physiological nor HF diet conditions revealed an effect of Wisp1 deficiency on whole-body glucose tolerance. However, male KO mice on a HF diet exhibited enhanced insulin sensitivity, lower insulin levels, and a marked reduction in adipose tissue inflammation, as evidenced by diminished macrophage infiltration and decreased pro-inflammatory cytokine expression in visceral fat. Additionally, beta cell mass expansion was attenuated in KO mice under HF diet, aligning with lower macrophage infiltration in islets. These findings suggest that improved insulin sensitivity in KO mice occurs independently of changes in glucose tolerance, likely due to mitigated adipose tissue inflammation. Thus, Wisp1 primarily modulates local adipose inflammatory responses, indirectly affecting islet adaptation to metabolic stress, rather than serving as a direct regulator of systemic glucose homeostasis.

## INTRODUCTION

The Cellular Communication Network (CCN) protein family consists of six matricellular proteins that are non-structural and secreted into the extracellular matrix (ECM), where they regulate intercellular signaling^1^. CCN proteins exhibit a conserved modular domain structure that allows for interactions with both structural components of the ECM, cell surface receptors and growth factors, thereby influencing a wide range of cellular events including differentiation, proliferation, adhesion, apoptosis or migration^2,3^. These proteins can also be found in significant amounts in circulation^4^, where they may exert endocrine effects on distant organs, in addition to their autocrine and paracrine functions.

CCN4/Wisp1 (Wnt1-inducible signaling pathway protein 1), was first identified in the late 1990s as a protein associated with cancer^5^. However, compared to other members of the CCN family, our understanding of Wisp1 remains relatively limited^6^. Despite this, a steadily growing body of literature implicates Wisp1 dysregulation and atypical expression profiles in a wide spectrum of pathophysiological conditions, including various cancers, fibrosis, arthritis, and metabolic disorders including obesity and diabetes^3,6,7^. In humans, Wisp1 is broadly expressed across multiple tissues, including skeletal muscle, lung, kidney, heart, liver, brain, ovaries, and adipose tissue^5,8^. Within these organs, it is expressed in distinct subsets of cells, for example, in fibroblasts, neurons, osteoblasts and adipocytes^6,9^.

Various lines of evidence suggest a link between Wisp1 and obesity. *Wisp1* expression is significantly elevated in visceral adipose tissue (VAT) in individuals with obesity and in mice fed a high fat diet^9,10^. In addition, serum Wisp1 levels are generally higher in human obesity, correlating with visceral adiposity and markers of systemic inflammation^9–13^. Further reinforcing its connection to inflammation, *Wisp1* expression has been associated with macrophage infiltration in non-obese glucose-tolerant individuals, and exogenous Wisp1 promotes a pro-inflammatory phenotype in macrophages *in vitro*^11^.

Wisp1 has also been implicated in the development of insulin resistance associated with obesity. Serum Wisp1 levels have been positively correlated with fasting insulin and negatively correlated with insulin sensitivity and circulating adiponectin^9^. *In vitro* studies have shown that Wisp1 inhibits insulin action in muscle and liver cells^10,14^. However, these findings contrast with other research suggesting that Wisp1 enhances insulin signaling in adipocytes, neurons, and pancreatic islets^15–17^.

In our previous work, we demonstrated that serum levels of Wisp1 are elevated during postnatal development in mice and in children compared to adults, likely reflecting increased production by growing bones^17^. Furthermore, we showed that Wisp1 is not endogenously expressed in pancreatic islets; however, exogenous Wisp1 promoted beta cell proliferation^17^, supporting a potential role for Wisp1 in the regulation of beta cell mass.

Overall, current evidence suggests that Wisp1 is linked to obesity, adipose tissue inflammation, insulin resistance and pancreatic beta cell mass, highlighting its potential role in the pathogenesis of type 2 diabetes (T2DM). Nevertheless, the precise nature of this relationship remains unclear. While elevated Wisp1 levels have been reported in gestational diabetes^18^ and T2DM^12,19^, other studies have found no significant association between circulating Wisp1 levels and glycemic status in obese or diabetic individuals^10–12^. These discrepancies may reflect differences in study design, population size and characteristics, or analytical methods.

Given the presence of conflicting data, primarily derived from correlation analysis in cross-sectional human studies or *in vitro* findings, we aimed to investigate the causal relationships underlying these processes using an *in vivo* preclinical model. To this end, we employed a systemic Wisp1 knockout (Wisp1 KO) mouse model to explore the role of Wisp1 in regulating adult glucose homeostasis under both physiological conditions and in the context of obesity-induced metabolic stress. Our study addresses a critical gap in the field, as preclinical models providing comprehensive metabolic analyses of Wisp1 deficiency in obesity and related disorders remain lacking.

## RESULTS

### Glucose homeostasis in Wisp1 KO mice under physiological conditions

To assess the role of Wisp1 in glucose homeostasis, we first performed intraperitoneal glucose tolerance tests on male and female 11-week-old Wisp1 KO mice maintained on a standard chow diet (SC, henceforth). KO mice exhibited similar body weight, fasting glucose levels, and glucose tolerance compared to wild-type (WT) controls in males and females (Fig. 1A–C). Likewise, fasting insulin levels and insulin tolerance tests showed no significant differences between genotypes in both sexes (Fig. 1D,E). Insulin secretion in response to glucose was comparable between WT and KO mice *in vivo* (Fig. 1F), and *in vitro* glucose-stimulated insulin release from isolated islets also showed no differences (Fig. S1). Morphometric analysis of the pancreas revealed comparable beta, alpha, and delta cell mass across genotypes in both sexes (Fig. 1G,H). In 50-week-old male mice, no significant differences in glucose homeostasis were observed between KO and WT groups, thus indicating that aging does not reveal any latent effects of Wisp1 deficiency on glucose regulation (Fig.S2). Together, these data indicate that, under physiological conditions, *Wisp1* is dispensable for maintaining normal glucose homeostasis in the adult.

**Figure 1.**
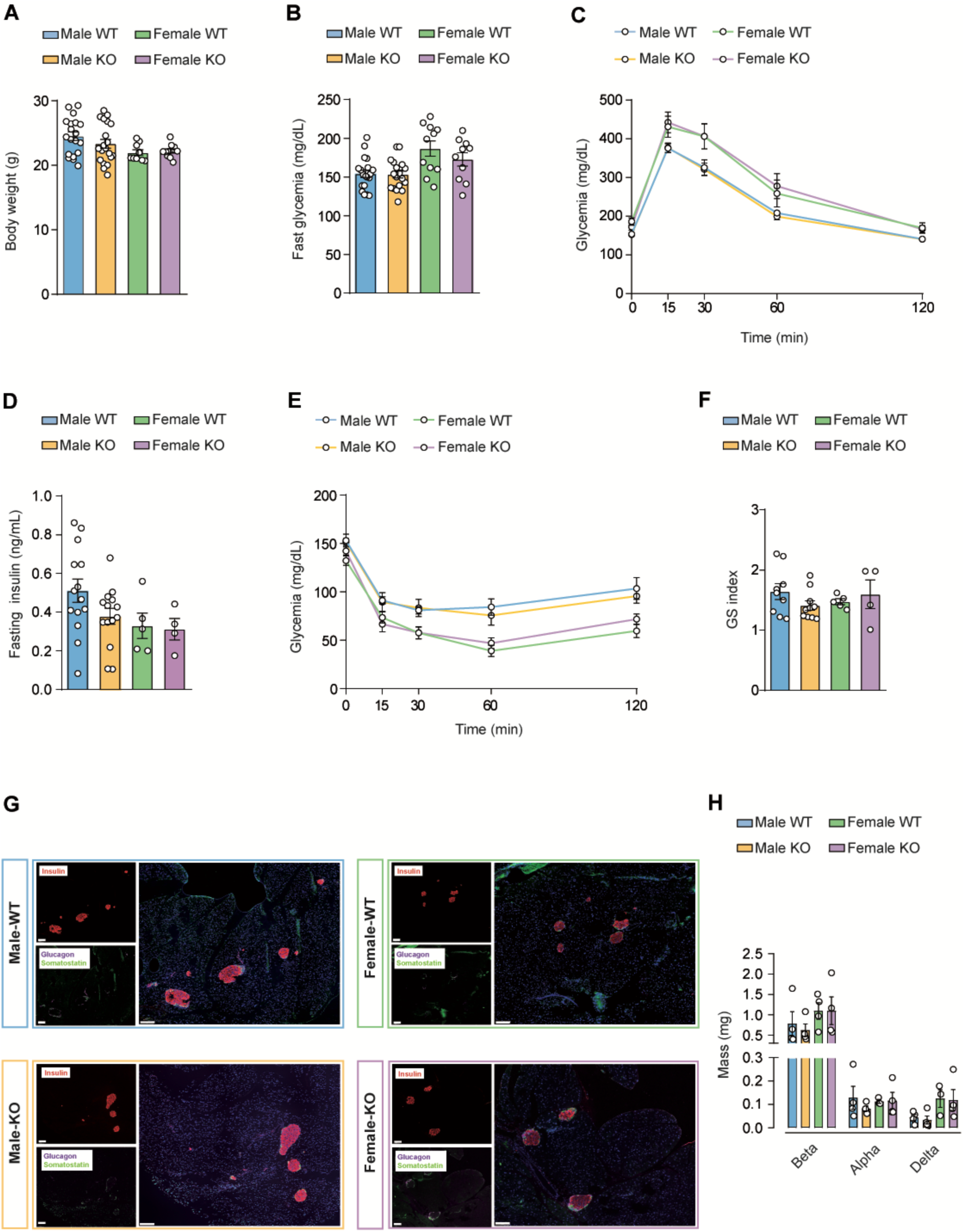
Glucose homeostasis in adult male and female Wisp1 KO mice under physiological conditions. **A.** Body weight of 11-week-old male and female Wisp1 WT and KO mice. Data are presented as mean±SEM for n=10-20 mice, shown as individual points. Comparisons were made using two-tailed Student’s *t* test within each sex. **B.** Five-hour fasting glycemia of 11-week-old male and female Wisp1 WT and KO mice. Data are presented as mean±SEM for n=10-20 mice, shown as individual points. Comparisons were made using two-tailed Student’s *t* test within each sex. **C.** ipGTT results in 11-week-old male and female Wisp1 WT and KO mice. Data are shown as mean±SEM for n=8-18 mice. Comparisons were made using a two-way ANOVA within each sex. **D.** Five-hour fasting plasma insulin levels of 11-week-old male and female Wisp1 WT and KO mice. Data are presented as mean±SEM for n=4-14 mice, shown as individual points. Comparisons were made using two-tailed Student’s *t* test within each sex. **E.** ipITT results in male and female Wisp1 WT and KO mice at 11 weeks of age. Data are shown as mean±SEM for n=10-15 mice. Comparisons were made using a two-way ANOVA within each sex. **F.** *In vivo* glucose-stimulated insulin index (GS index), calculated as insulin levels 15 minutes after glucose injection relative to basal insulin levels following a five-hour fasting in11-week-old male and female Wisp1 WT and KO mice. Data are presented as mean±SEM for n=4-10 mice, shown as individual points. Comparisons were made using two-tailed Student’s *t* test within each sex. **G.** Representative immunofluorescence images of fixed pancreatic sections from 11-week-old male and female Wisp1 WT and KO mice stained for insulin (red), glucagon (purple), and somatostatin (green). Nuclei were stained using Hoechst (blue). Scale bar: 100μm **H.** Beta-, alpha- and delta-cell mass was assessed in fixed pancreatic sections of 11-week-old male and female Wisp1 WT and KO mice using morphometric quantification. Data are presented as mean±SEM from n=3-4 mice, shown as individual points. Comparisons were made using two-tailed Student’s *t* test within each sex and for each hormone.

### Glucose homeostasis in Wisp1 KO mice under diet-induced obesity conditions

Given the reported association of Wisp1 and human obesity, we reasoned that its functional role might only become evident under metabolic stress associated with this condition. Therefore, we studied the impact of Wisp1 deficiency in the context of diet-induced obesity. Six-week-old male KO and WT mice were fed either a high fat (HF) or a SC diet for 20 weeks. Body weight gain in KO mice was comparable to that of WT controls (Fig. 2A), and HF diet-induced changes in body composition, including VAT (represented by epididymal white adipose tissue or eWAT) and subcutaneous adipose tissue (scWAT), did not differ significantly between genotypes (Fig. 2B). Similarly, pancreas and liver weights showed no significant genotype-dependent differences following a HF diet (Fig. S3). No significant differences were observed in plasma triglycerides or free fatty acids (FFAs) under fed conditions across groups (Fig. 2C,D). Yet, in contrast to WT mice, KO mice maintained a significant fed-to-fasted increase in plasma FFA levels despite HF diet feeding (Fig. 2D).

**Figure 2.**
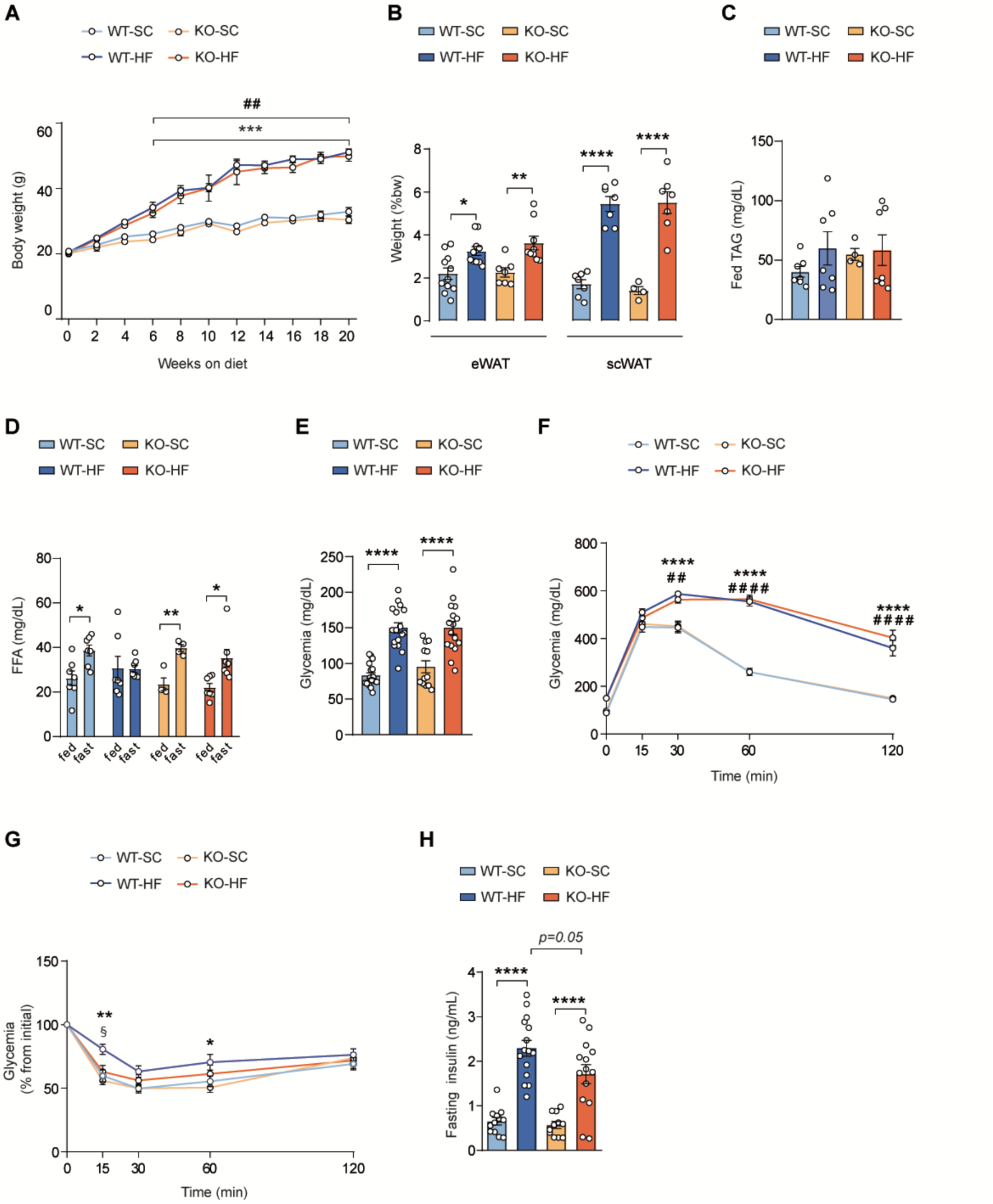
Glucose homeostasis in male Wisp1 KO mice under HF diet conditions. **A.** Body weight was monitored in male Wisp1 WT and KO mice fed either a HF or a SC diet for 20 weeks. Data are shown as mean±SEM from n=8-17 mice. ***p<0.001 (WT-SC *vs* WT-HF) ##p<0.01 (KO-SC *vs* KO-HF). Comparisons were made using a two-way ANOVA. **B.** Comparison of the relative eWAT and scWAT weights normalized to total body weight in male Wisp1 WT and KO mice fed either a HF or a SC diet for 20 weeks. Data are shown as mean±SEM from n=7-11 mice, shown by individual points. *p<0.05, **p<0.01, ****p<0.0001. Comparisons were made using a one-way ANOVA within each depot. **C.** Fed plasma triglyceride (TAG) levels in male Wisp1 WT and KO mice fed either a HF or a SC diet for 20 weeks. Data are shown as mean±SEM from n=4-7 mice, shown as individual points. Comparisons were made using a one-way ANOVA. **D.** Fed and fasted plasma free fatty acid (FFA) levels in male Wisp1 WT and KO mice fed either a HF or a SC diet for 20 weeks. Data are shown as mean±SEM from n=4-7 mice, shown as individual points. *p<0.05, **p<0.01. Comparisons were made using multiple two-tailed Student’s *t* test. **E.** Overnight fasting glycemia in male Wisp1 WT and KO mice fed HF or SC diets for 19 weeks. Data are presented as mean±SEM from n=12-18 mice, shown as individual points. ****p<0.0001. Comparisons were made using one-way ANOVA. **F.** ipGTT results in male Wisp1 WT and KO mice fed either a HF or a SC diet for 19 weeks. Data are shown as mean±SEM from n=12-18 mice. ##p<0.01 (KO-SC *vs* KO-HF), ****p<0.0001 (WT-SC *vs* WT-HF), #### p<0.0001 (KO-SC *vs* KO-HF). Comparisons were made using two-way ANOVA. **G.** ipITT results in male Wisp1 WT and KO mice fed either a HF or a SC diet for 18 weeks. Results are expressed as the percentage of variation from initial glucose concentration (100%). Data are presented as the mean±SEM for n=12-18 mice. § p<0.05 (WT-HF *vs* KO-HF), *p<0.05 (WT-SC *vs* WT-HF) **p<0.01 (WT-SC *vs* WT-HF). Comparisons were made using two-way ANOVA. **H.** Plasma insulin levels after an overnight fast in male Wisp1 WT and KO mice fed either a HF or a SC diet for 19 weeks. Data are presented as mean±SEM from n=11-15 mice, shown as individual points. ****p<0.0001. Comparisons were made using one-way ANOVA.

Both KO and WT mice developed comparable levels of fasting hyperglycemia and glucose intolerance (Fig. 2E,F). However, KO mice demonstrated significantly improved insulin tolerance relative to WT mice, with responses closely resembling those observed in SC-fed KO and WT controls (Fig. 2G). This enhanced insulin sensitivity in HF-fed KO mice was further supported by lower fasting insulin levels compared to their WT counterparts (Fig. 2H).

We applied the same HF diet regimen to female Wisp1 KO mice and confirmed that Wisp1 deficiency did not confer any advantage in glucose handling under HF diet conditions. Both KO and WT females exhibited comparable increases in body weight, body composition, fasting glucose levels, and glucose intolerance (Fig. S4A-D). Consistent with previous reports^20,21^, and unlike their male counterparts, female mice were resistant to HF-induced reduction in insulin sensitivity and did not develop fasting hyperinsulinemia, and Wisp1 absence did not modify this phenotype (Fig. S4E,F).

Collectively, these findings show that Wisp1 deficiency confers a protective effect on insulin sensitivity in male mice in the context of diet-induced obesity. However, this improvement does not translate into enhanced whole-body glucose tolerance.

### Systemic Wisp1 levels and adipose-specific *Wisp1* gene expression changes in diet induced obesity

We next sought to identify the mechanism/s responsible for the ameliorated insulin resistance observed in HF diet-fed male Wisp1 KO mice. We began by characterizing the temporal profile of circulating Wisp1 levels in WT mice subjected to either SC or HF diets over a 24-week period. We found that plasma Wisp1 concentrations did not differ significantly between SC and HF-fed groups at the time points analyzed; however, levels gradually declined over time under both dietary conditions, likely reflecting the age-related reduction previously reported^17^ (Fig. 3A). Given the absence of systemic Wisp1 changes in response to the HF diet, we examined tissue-specific *Wisp1* gene expression in WT mice to identify potential sites of functional relevance. Consistent with earlier reports in humans and murine models^9,10,15^, we found that basal *Wisp1* mRNA levels were higher in eWAT compared to scWAT. Notably, its expression was significantly upregulated under HF diet in eWAT. By contrast, *Wisp1* expression remained unchanged in scWAT and liver, and was marginally expressed or undetectable in skeletal muscle and islets under both SC and HF diet conditions (Fig. 3B).

**Figure 3.**
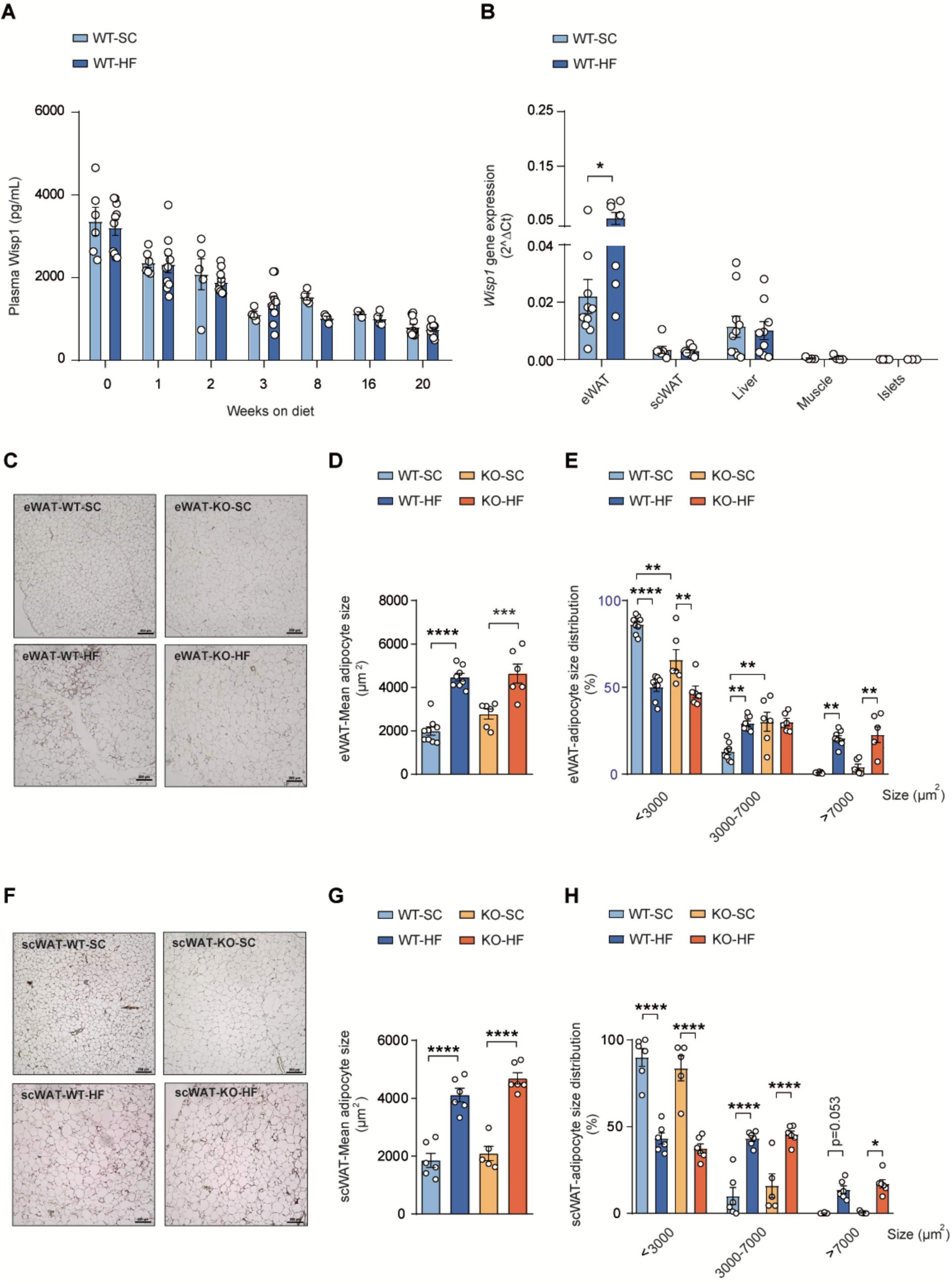
Patterns of *Wisp1* gene expression and circulating levels during HF feeding, and the impact of Wisp1 deficiency on adipose tissue morphology. **A.** Plasma Wisp1 levels in male WT mice fed either a HF or a SC diet for 20 weeks. Data are presented as mean±SEM for n=4-11 mice shown as individual points. Comparisons were made using two-Way ANOVA. **B.** *Wisp1* gene expression in the indicated tissues harvested from male WT mice fed either a HF or a SC diet for 20 weeks. Values are expressed relative to the average of the housekeeping genes *Rplp0* and *Tbp*. Data are presented as mean±SEM from n=4-9 mice shown as individual points. *p<0.05. Comparisons were made using two-tailed Student’s *t* test. **C.** Representative hematoxilin & eosin staining of eWAT from male Wisp1 WT and KO mice fed either a HF or a SC diet for 20 weeks. Scale bar 200μm. **D.** Mean adipocyte size in eWAT from male Wisp1 WT and KO mice fed either a HF or a SC diet for 20 weeks. Data are shown as mean±SEM from n=6-9. ***p<0.001, ****p<0.0001. Comparisons were made using a one-way ANOVA. **E.** Adipocyte size distribution in eWAT from male Wisp1 WT and KO mice fed either a HF or a SC diet for 20 weeks. Data are shown as mean±SEM from n=5-9. **p<0.01, ****p<0.0001. Comparisons were made using a one-way ANOVA. **F.** Representative hematoxilin & eosin staining of scWAT from male Wisp1 WT and KO mice fed HF or SC diets for 20 weeks. Scale bar 200μm. **G.** Mean adipocyte size in scWAT from male Wisp1 WT and KO mice fed either a HF or a SC diet for 20 weeks. Data are shown as mean±SEM from n=6-9. ***p<0.001, ****p<0.0001. Comparisons were made using a one-way ANOVA. **H.** Adipocyte size distribution in scWAT from male Wisp1 WT and KO mice fed either a HF or a SC diet for 20 weeks. Data are shown as mean±SEM from n=5-6. *p<0.05 ****p<0.0001. Comparisons were made using a one-way ANOVA.

These findings suggest that Wisp1 likely acts through local autocrine or paracrine mechanisms rather than the systemic circulation, with visceral adipose tissue emerging as a key site of action during diet-induced obesity.

### Adipose tissue morphology and function-related gene expression in Wisp1 KO mice

Based on the above results, we next focused on the effects of Wisp1 deficiency in eWAT. As an initial step, we evaluated whether other CCN family members might compensate for the loss of Wisp1 in KO mice. Expression levels of *Ccn1(Cyr61), Ccn2*(*Ctgf*), as well as *Ccn5* (*Wisp2)*, previously linked to insulin sensitivity^22,23^, did not differ significantly between genotypes under either SC or HF diets in either eWAT or scWAT (Fig. S5A).

Histomorphometric analysis of eWAT revealed that a HF diet increased the mean adipocyte size in both KO and WT mice. Although no significant size differences were observed between genotypes under HF diet, KO mice showed a slightly smaller increase (Fig. 3C,D). Under SC diet, KO mice displayed fewer small adipocytes and more medium-sized ones, likely accounting for their higher mean size. However, this distribution shift disappeared with HF feeding (Fig. 3E). On the other hand, in scWAT, neither mean adipocyte size nor size distribution differed significantly between genotypes under SC or HF diet conditions (Fig. 3F–H). In addition, gene expression analysis of selected adipocyte differentiation, lipolysis and lipogenic markers showed no significant differences between KO and WT mice fed a HF diet in both depots (Fig. S5B).

Together, these findings argue that Wisp1 deficiency does not result in overt alterations in adipose tissue morphology or in the expression of key functional genes during diet-induced obesity.

### Effects of diet-induced obesity on selected adipokines in Wisp1 KO mice

We next examined gene expression of adipocyte-derived proteins (adipokines) that have been associated with the maintenance of whole-body glucose balance and systemic insulin sensitivity^24–27^, including leptin (*Lep*), adiponectin (*Adipo*q), adipsin (*Cfd*) and resistin (*Retn*).

In eWAT, *Lep* mRNA levels remained unchanged following the HF diet in both genotypes (Fig. 4A). In contrast, *Adipoq*, *Cfd*, and *Retn* expression decreased in response to the HF diet in both KO and WT mice (Fig. 4A). Notably, under the SC diet, *Adipoq* expression was higher in KO mice compared to WT mice but this difference was abolished under HF conditions as expression declined in both genotypes. In scWAT, the HF diet markedly increased *Lep* expression in both WT and KO mice; however, this induction was attenuated in KO mice, resulting in lower *Lep* mRNA levels compared to WT under HF conditions (Fig. 4B). Conversely, *Adipoq* and *Retn* expression did not vary across diets or genotypes, while *Cfd* expression decreased in response to the HF diet similarly in both genotypes (Fig. 4B). These findings reveal a selective decrease of *Lep* expression in scWAT of KO mice fed a HF diet compared to WT mice, with no differences detected for this adipokine in eWAT.

**Figure 4.**
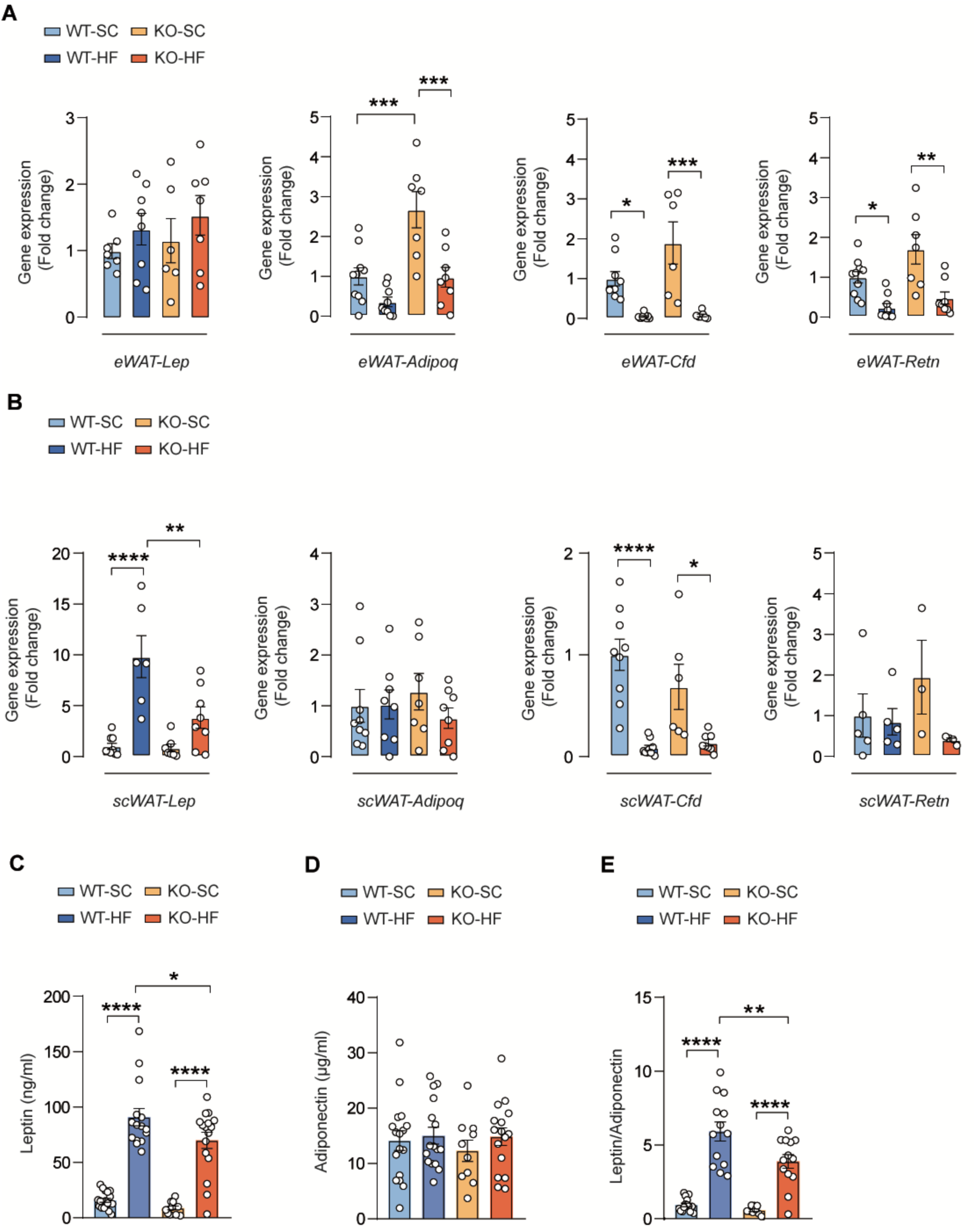
Adipokine gene expression and circulating levels in male Wisp1 KO mice under HF diet conditions. **A.** Gene expression of the indicated adipokines in eWAT from male Wisp1 WT and KO mice fed either a HF or a SC diet for 20 weeks. Expression was assessed by qPCR, using *Rplp0* and *Tbp* as housekeeping genes. Results are expressed relative to WT-SC, given the value of 1. Data are shown as mean±SEM from n=6-9. *p<0.05, **p<0.01, ***p<0.001. Comparisons were made using one-way ANOVA. **B.** Gene expression of the indicated adipokines in scWAT from male Wisp1 WT and KO mice fed either a HF or a SC diet for 20 weeks. Expression was assessed by qPCR, using *Rplp0* and *Tbp* as housekeeping genes. Results are expressed relative to WT-SC, given the value of 1. Data are shown as mean±SEM from n=3-9. *p<0.05, **p<0.01, ****p<0.0001. Comparisons were made using one-way ANOVA. **C.** Serum leptin levels in male Wisp1 WT and KO mice fed either a HF or a SC diet for 20 weeks. Data are presented as mean±SEM for n=11-18 mice shown as individual points. *p<0.05, ****p<0.0001. Comparisons were made using a one-way ANOVA. **D.** Serum adiponectin levels in male Wisp1 WT and KO mice fed either a HF or a SC diet for 20 weeks. Data are presented as mean±SEM for n=10-17 mice shown as individual points. Comparisons were made using a one-way ANOVA. **E.** Leptin to adiponectin ratio in male Wisp1 WT and KO mice fed either a HF or a SC diet for 20 weeks. Data are presented as mean±SEM for n=10-17 mice shown as individual points. **p<0.01, ****p<0.0001. Comparisons were made using a one-way ANOVA.

Finally, we assessed whether these gene expression changes were reflected in plasma. Consistent with the reduction of *Lep* mRNA in scWAT, KO mice displayed approximately 28% lower plasma leptin compared to WT under HF diet (Fig. 4C). On the other hand, in agreement with the lack of changes in *Adipoq* expression, plasma adiponectin levels remained similar across genotypes and diets (Fig. 4D). Consequently, the leptin:adiponectin ratio, used as biomarker of insulin resistance^28^, was significantly lower in KO mice relative to WT mice under HF conditions (Fig. 4E).

In summary, Wisp1 KO mice on a HF diet exhibit reduced circulating leptin, likely due to decreased *Lep* expression in scWAT. Since Wisp1 expression is unchanged in this fat depot and insulin is a known regulator of leptin production^29,30^, the reduction in leptin observed in KO mice under HF conditions may be secondary to lower insulin levels rather than a primary alteration in leptin regulation. These findings suggest that Wisp1 deficiency may influence other factors affecting insulin sensitivity and insulin levels, warranting further investigation in VAT to clarify the underlying mechanisms.

### Effects of diet-induced obesity on adipose tissue fibrosis in Wisp1 KO mice

Fibrotic remodeling of adipose tissue contributes to obesity-associated metabolic dysfunction, such as insulin resistance^31^. Given Wisp1’s established role in fibrosis in other tissues^8,32,33^, we next investigated its involvement in VAT fibrosis in Wisp1 KO mice. Masson’s Trichrome staining showed no major fibrotic differences in eWAT between groups (Fig. S6A). We also assessed the expression of ECM-related genes, including various collagens and integrins, but observed no significant differences under either dietary condition, except for reduced *Tgfbr1* expression in KO compared to WT, which reached significance only under HF diet conditions (Fig. S6B).

In scWAT, Masson’s Trichrome staining did not reveal fibrotic areas in either KO or WT tissue under SC or HF diet conditions (Fig. S7A). Gene expression analysis showed moderate differences in *Fn1* and *Tpm4* between KO and WT under HF diet, along with a blunted upregulation of *ItgaV* in KO mice in response to HF diet (Fig. S7B).

Together, these findings suggest that Wisp1 deficiency induces changes in a limited set of fibrosis-related genes in adipose tissue, including a notably reduced expression of *Tgbr1* in eWAT, a key driver of fibrotic signaling. However, these transcriptional alterations do not translate into detectable fibrotic changes in adipose tissue under the tested conditions, indicating that fibrosis is unlikely to be the primary mechanism underlying improved insulin tolerance in this murine model.

### Effects of diet-induced obesity on adipose tissue inflammation in Wisp1 KO mice

In the context of obesity, VAT also undergoes extensive inflammatory remodeling, impairing its metabolic function and contributing to systemic insulin resistance ^34,35^. As Wisp1 has been implicated in pro-inflammatory signaling, we assessed inflammatory responses in Wisp1 KO mice under HF diet conditions. To address this aim, we assessed macrophage infiltration by F4/80 staining and observed a pronounced reduction in F4/80-positive staining in eWAT from KO mice compared with WT controls under HF diet conditions (Fig. 5A). In contrast, no significant differences were detected between KO and WT mice in scWAT (Fig. S8A). Whole mount 3D tissue imaging using Cd68 as a pan-macrophage marker also showed decreased macrophage presence in eWAT of KO mice (Video S1) relative to WT (Video S2).

**Figure 5.**
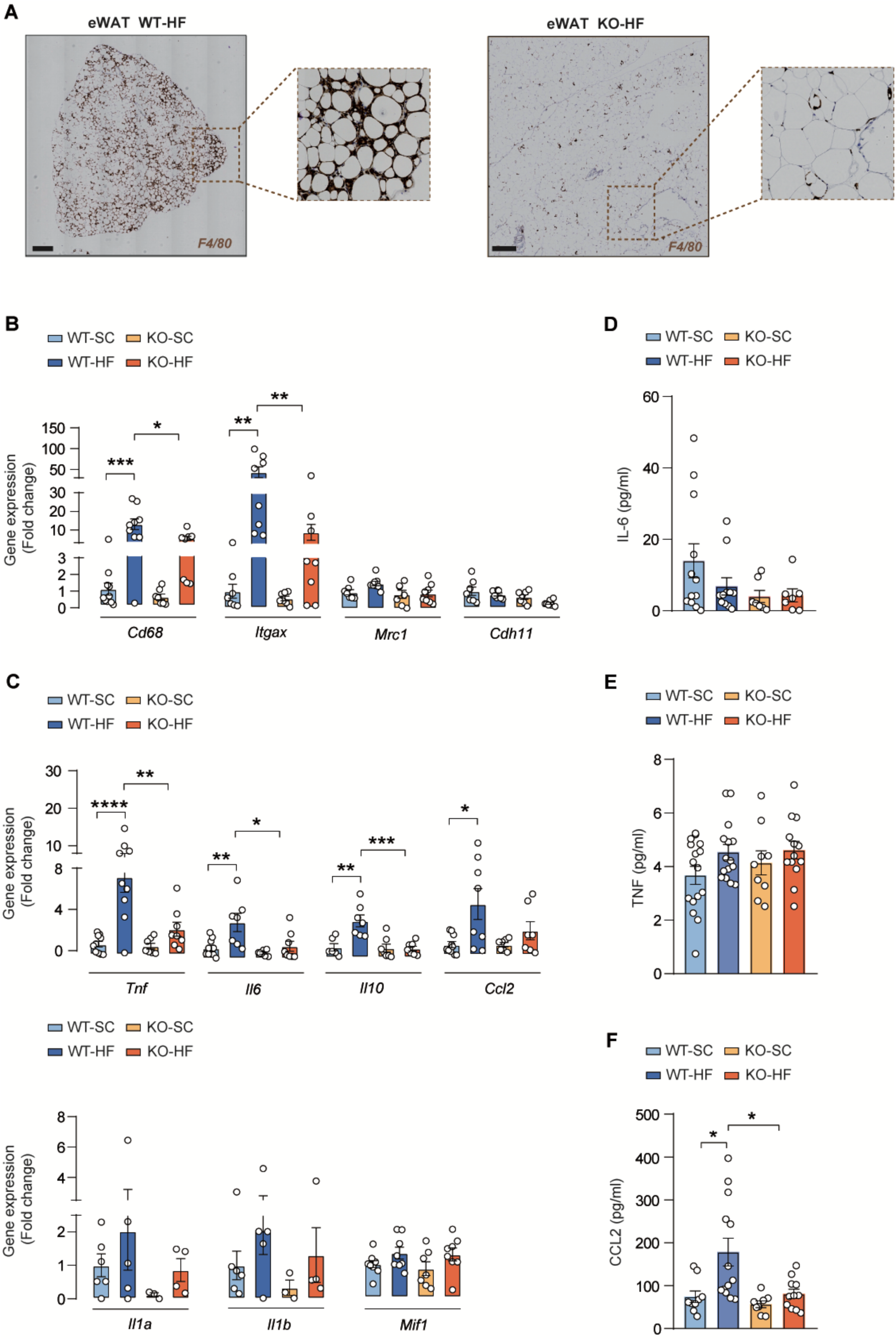
Assessment of eWAT inflammation in male Wisp1 KO mice under HF diet conditions. **A.** Representative immunohistochemistry images showing positive F4/80 staining in eWAT from male Wisp1 WT and KO mice fed a HF diet for 20 weeks. Scale bar 500μm. **B.** Gene expression of macrophage-related genes in male Wisp1 WT and KO mice fed either a HF or a SC diet for 20 weeks. Expression was assessed by qPCR, using *Rplp0* and *Tbp* as housekeeping genes. Results are expressed relative to WT-SC, given the value of 1. Data are presented as mean±SEM for n=7-10 mice. *p<0.05, **p<0.01, ***p<0.001. Comparisons were made using a one-way ANOVA. **C.** Gene expression of inflammation-related genes in male Wisp1 WT and KO mice fed either a HF or a SC diet for 20 weeks. Expression was assessed by qPCR, using *Rplp0* and *Tbp* as housekeeping genes. Results are expressed relative to WT-SC, given the value of 1. *p<0.05, **p<0.01, ***p<0.001, ***p<0.0001. Comparisons were made using a one-way ANOVA. **D.** Serum IL-6 levels in male Wisp1 WT and KO mice fed either a HF or a SC diet for 20 weeks. Data are presented as mean±SEM for n=7-12 mice. Comparisons were made using a one-way ANOVA. **E.** Serum TNF levels in male Wisp1 WT and KO mice fed either a HF or a SC diet for 20 weeks. Data are presented as mean±SEM for n=9-16 mice. Comparisons were made using a one-way ANOVA.**F.** Serum CCL2 levels in male Wisp1 WT and KO mice fed either a HF or a SC diet for 20 weeks. Data are presented as mean±SEM from n=6-11 mice. *p<0.05. Comparisons were made using a one-way ANOVA.

To further support these findings, we analysed the expression of genes linked to macrophage presence and inflammatory signaling in eWAT. Consistent with the imaging data, *Cd68* expression was significantly reduced in KO mice (Fig. 5B). Likewise, *Itgax*, which encodes Cd11c, a marker of pro-inflammatory M1-like macrophages and dendritic cells, was markedly decreased in eWAT from KO mice. In contrast, expression levels of *Mrc1* (Cd106, an M2 macrophage marker) and *Cdh11* (a marker of activated fibroblasts) remained unchanged (Fig. 5B).

Additionally, KO mice also exhibited significantly lower expression, or a blunted HF diet induced upregulation, of pro-inflammatory cytokine genes including *Tnf, Il6, Il10*, and the chemokine *Ccl2* (a key mediator of macrophage recruitment). In contrast, expression levels of *Il1a, Il1b* or *Mif1* remained unchanged (Fig. 5C). To determine whether these changes were specific to this adipose depot, we examined the mRNA levels of some of these genes in scWAT. Compared to eWAT, scWAT exhibited fewer inflammation-related gene expression changes in response to HF diet, and when changes were present, the fold increases were markedly lower. Nonetheless, as in eWAT, the inductions of the *Itgax* and *Ccl2* genes under HF diet were blunted in KO mice compared to WT controls in scWAT (Fig. S8B).

To assess the systemic impact of these gene expression changes, we measured circulating levels of inflammatory cytokines. No significant differences were observed in IL-6 or TNF concentrations between Wisp1 KO and WT mice upon HF feeding (Fig. 5D,E). By contrast, consistent with the transcriptional data, HF-fed KO mice exhibited a blunted increase in circulating CCL2 levels as compared to WT mice (Fig. 5F).

In conclusion, *Wisp1* deficiency mitigates HF diet-induced eWAT inflammation by limiting macrophage infiltration and suppressing pro-inflammatory gene expression. Inflammatory gene expression is also reduced, although less pronounced, in scWAT. These changes likely contribute to the improved systemic insulin tolerance observed in KO mice, with reduced HF-diet induced circulating Mcp-1 levels possibly representing one underlying mechanism.

### Effects of diet-induced obesity on beta cell mass and function in Wisp1 KO mice

We next asked why whole-body glucose tolerance remained unaltered in HFD-fed Wisp1 KO mice despite improved insulin sensitivity. We postulated that Wisp1 deficiency might impair adaptive beta cell expansion, thereby limiting glucose tolerance improvements. To test this, we performed morphometric analysis of pancreatic tissue to quantify beta cell mass. In response to a HF diet, WT mice exhibited a robust 5.7-fold increase in beta cell mass (p < 0.01), whereas KO mice showed a modest, non-significant 1.8-fold increase (Fig. 6A,B). This expansion was accompanied in WT mice by a shift in islet size distribution, consisting of a reduction in small islets and an increase in large islets, indicative of compensatory beta cell growth. In contrast, KO mice maintained a stable islet size profile under HF diet conditions (Fig. 6C). HF feeding did not affect alpha cell mass or glucagon levels, indicating that effects are restricted to the beta-cell compartment (Fig. S9). To assess beta cell functionality, we performed *in vivo* and *in vitro* glucose-stimulated insulin secretion assays. These experiments revealed no significant differences in glucose responsiveness between Wisp1 KO and WT groups either *in vivo* (Fig. 6D) or *ex vivo* using isolated islets (Fig. S10), indicating that Wisp1 deficiency does not substantially change beta cell function under HF diet conditions. Collectively, these findings suggest that the absence of adaptive beta cell expansion in Wisp1 KO mice may underlie their inability to improve glucose tolerance despite exhibiting enhanced insulin sensitivity.

**Figure 6.**
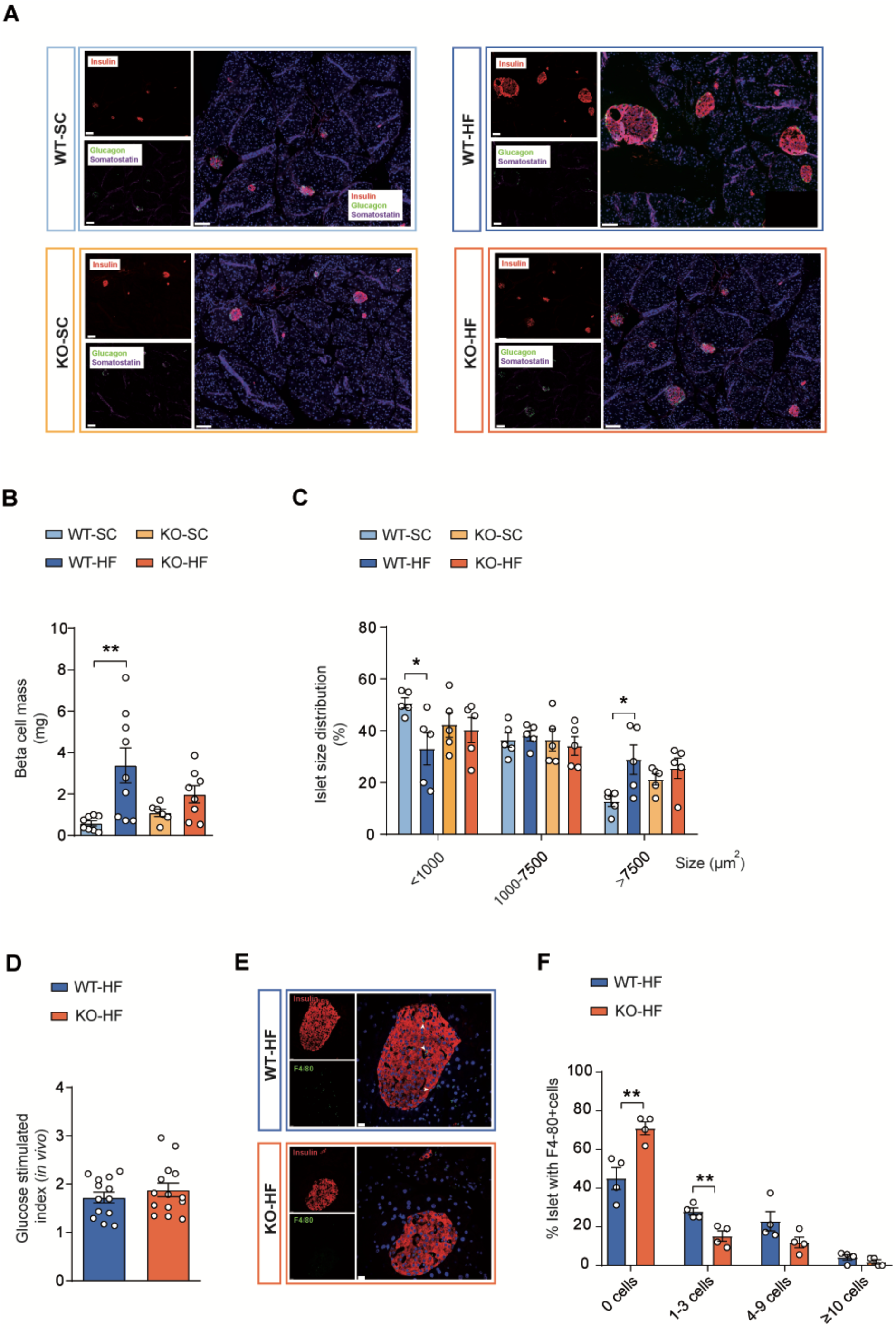
Characterization of the beta cell compartment in male Wisp1 KO mice under HF conditions. **A.** Representative immunofluorescence images of formalin-fixed paraffin-embedded pancreatic sections from male Wisp1 WT and KO mice fed either a HF or a SC diet for 20 weeks. Sections were stained for insulin (red), glucagon (green), and somatostatin (purple). Nuclei were stained using Hoechst (blue). Scale bar: 100μm **B.** Morphometric quantification of beta-cell mass in fixed pancreatic sections from male Wisp1 WT and KO mice fed either a HF or a SC diet for 20 weeks. Data are shown as mean±SEM from n=6-9 mice. **p<0.01. Comparisons were made using a one-way ANOVA. **C.** Islet size distribution in male Wisp1 WT and KO mice fed either a HF or a SC diet for 20 weeks. Results represent percentage of total islet number for each size category. Data are presented as mean±SEM for n=5. *p<0.05. Comparisons were made using a one-way ANOVA. **D.** *In vivo* glucose-stimulated index, calculated as insulin levels 15 minutes after glucose injection relative to basal insulin levels following an overnight fast in male Wisp1 WT and KO mice fed a high fat diet for 19 weeks. Data are shown as mean±SEM from n=13-14. Comparison was made using two-tailed Student’s *t* test. **E.** Representative immunofluorescence images of pancreatic islets in formalin-fixed paraffin-embedded pancreatic sections from male Wisp1 WT and KO mice fed a high fat diet for 20 weeks, stained for insulin (red) and F4/80 (green). Nuclei were stained using Hoechst (blue). Scale bar: 100μm. **F.** Quantification of F4-80-positive cells within the pancreatic islets of male Wisp1 WT and KO mice fed a high fat diet for 20 weeks. Results are represented as the percentage of islets with the indicated number of F4/80-positive cells. Data are expressed as mean±SEM from n=4. **p<0.01. Comparisons were made using multiple Student’s t-test.

Finally, we investigated whether reduced adipose tissue inflammation might be linked to impaired beta cell expansion in Wisp1 KO mice under HF conditions. Macrophage signaling has been implicated in promoting beta cell proliferation in several beta cell regeneration models^36–38^ as well as in obesity^39^. Given the lower circulating CCL2 levels detected in HF-fed Wisp1 KO mice and its known role in promoting monocyte recruitment to tissues^40,41^, we assessed macrophage presence in islets from KO and WT mice on a HF diet. F4/80 immunostaining was performed to quantify intra-islet macrophages. Strikingly, KO pancreases exhibited a higher proportion of islets devoid of macrophages compared to WT (Fig. 6E,F), suggestive of reduced local immune cell infiltration. This diminished presence of inflammatory cells within islets, cells previously shown to support compensatory beta cell mass expansion, may underlie the limited beta cell growth observed in Wisp1 KO mice under obesity.

## DISCUSSION

This study presents the first *in vivo* loss-of-function analysis of *Wisp1* in the context of HF diet-induced obesity. Despite the impaired glucose tolerance typically associated with obesity, *Wisp1* deficiency neither worsens nor improves this condition in either male or female mice, as glucose clearance remains unaffected. However, in males, insulin sensitivity is enhanced, in association with lower circulating insulin levels and reduced pancreatic beta cell mass. These metabolic changes are accompanied by decreased inflammation in visceral adipose tissue, suggesting that Wisp1 contributes to insulin resistance and beta cell expansion through inflammatory pathways. Together, these findings reveal that Wisp1 plays a sex-specific role primarily as a regulator of adipose tissue inflammation and islet adaptation in obesity, rather than by acting as a direct modulator of systemic glucose metabolism.

Since its identification as an adipokine, Wisp1 has attracted attention as a potential biomarker for obesity and glucose homeostasis. Several studies in human cohorts, including both adults and children, have reported positive association between circulating WISP1 levels, BMI and total adiposity^10,12,13^. However, this association was weak or not seen in other studies^9,11,19^. For instance, reduction of body fat following weight loss did not alter circulating WISP1 levels in males and showed only a modest trend in females ^9^. These discrepancies could, in part, be attributed to difficulties in the detection of plasma WISP1 levels in adult individuals, which often fall below the detection limits of currently available assays^11^. In our study, while we detected increased *Wisp1* expression in VAT, which aligns with previous findings in mice and humans^9,15^, this increase did not translate into significant changes in circulating Wisp1 levels following a HF diet exposure, reinforcing the notion that Wisp1 may not be a reliable systemic biomarker for obesity.

Studies investigating the relationship between circulating Wisp1 levels and glycemic control in humans have also yielded conflicting results. While some reports suggest that systemic WISP1 concentrations vary with glycemic status^13,18^, others have found no consistent association between circulating WISP1 levels and glucose tolerance or T2DM^10–12^. Our current results align with these latter studies, as *Wisp1* deficiency does not appear to affect glycemia or systemic glucose tolerance under either normal physiological conditions or in the context of obesity, in both male and female mice. These findings reinforce the notion that Wisp1 may play a limited or non-essential role in the regulation of glucose homeostasis.

Yet, in response to HF diet-feeding, male Wisp1 KO mice exhibit lower fasting insulin levels and improved insulin tolerance as compared to their WT counterparts, reflecting enhanced insulin sensitivity despite no significant change in glucose tolerance. The improved insulin tolerance observed in males could be consistent with human studies reporting associations between WISP1 levels and markers of insulin sensitivity^9,10,18^, and is also in line with *in vitro* findings suggesting that Wisp1 protein can impair insulin signaling in hepatocytes and skeletal muscle^10,14^. In our model, however, circulating Wisp1 levels and its expression in muscle and liver were not altered by the HF diet, indicating that any effects of Wisp1 on these organs are likely indirect.

We identify VAT as a principal site of Wisp1 expression in obesity, suggesting that this tissue plays a central role in mediating Wisp1’s influence on whole-body metabolism. While we did not detect morphological changes or fibrosis in eWAT of Wisp1 KO mice fed a HF diet, our work provides the first *in vivo* causal evidence that Wisp1 drives adipose tissue inflammation during obesity. Mechanistically, Wisp1 deficiency reduces macrophage infiltration and downregulates pro-inflammatory gene expression in eWAT of male mice under HF diet conditions. These findings align with previous human studies reporting that *WISP1* gene expression in VAT is positively correlated with immune cell infiltration in both VAT and SAT depots^9^. In contrast to humans, our murine model showed no differences in macrophage infiltration and only changes in a few inflammatory markers in scWAT, likely reflecting the inherent low immune baseline infiltration in this tissue^42,43^, which may limit the ability to detect of Wisp1-dependent effects. At the molecular level, Wisp1 has been demonstrated to activate pro-inflammatory cytokine gene expression in macrophages^9^, supporting a paracrine mechanism by which Wisp1 produced by VAT adipocytes recruits and activates inflammatory macrophages locally.

Given the established role of localized adipose tissue inflammation in promoting systemic inflammation and impairing insulin action, reduced inflammation in eWAT of male *Wisp1* KO mice may explain their improved insulin sensitivity. Previous studies have shown that circulating WISP1 levels and its expression in VAT are positively associated with inflammatory markers in humans, including HO-1, IL-8, IL-18 or hsCRP^9,10,13,19^. In male Wisp1 KO mice, we identified CCL2, a potent chemokine involved in monocyte recruitment, as a possible link between Wisp1 activity in VAT and systemic inflammation. HF diet-induced *Ccl2* gene activation in VAT was attenuated in Wisp1 KO mice, accompanied by lower systemic CCL2 levels. Interestingly, *CCL2* expression has been shown to correlate with *WISP1* expression in human VAT ^10^.

Despite improved insulin sensitivity, male Wisp1 KO mice fed a HF diet do not fully restore normal glucose homeostasis. Notably, these mice display a blunted beta cell expansion response, which may result from decreased insulin demand linked to reduced eWAT inflammation. Although Wisp1 was previously identified as a pro-proliferative factor for beta cells during postnatal development^17^, its lack of expression in islets and stable systemic levels under HF diet suggest an indirect role in beta cell adaptation during adult obesity. In our study, we detected reduced macrophage infiltration in islets from Wisp1 KO mice, that we associated with reduced adipose tissue inflammation and decreased circulating levels of inflammatory molecules, CCL2 among them. In fact, proliferation of resident macrophages and release of trophic factors such as PDGF have been implicated in beta cell expansion in response to a HF diet in mice^39^. Our findings establish a directional regulatory link between inflammation in adipose tissue and pancreatic islets mediated by Wisp1, a molecule specifically upregulated in adipose tissue under HF feeding. While Wisp1 appears to play a significant role, it is unlikely to be the sole mediator, as modest beta cell expansion was still observed in KO mice. The precise mechanisms driving beta cell growth in insulin-resistant states remain incompletely understood but likely involve a complex interplay of systemic and local signals.

In summary, our study highlights the complex interplay between inflammation and beta cell plasticity in obesity. Wisp1 deficiency, by reducing VAT inflammation, indirectly limits beta cell expansion, revealing a trade-off between improved insulin sensitivity and adaptive islet growth. These findings underscore the complexity of targeting inflammatory pathways in type 2 diabetes; while dampening inflammation may improve insulin sensitivity, it could also inadvertently impair beta cell adaptability, potentially limiting therapeutic benefit.

## MATERIAL AND METHODS

### Mice

Wisp1 KO mice^44^ were bred at the barrier animal facility of the University of Barcelona and maintained in a 12-h light/12-h dark cycle in temperature and humidity-controlled environment with free access to water and standard chow (SC) diet. Genotyping was performed with primers provided in Supplementary Table 1. For diet-induced obesity model, 6-week-old male and female Wisp1 KO mice were fed a high fat (HF) diet (60% fat calories, 20% protein calories and 20% carbohydrate calories; Research Diets #D12492) for 20 weeks. Intraperitoneal insulin tolerance tests (ipITT) and glucose tolerance (ipGTT) tests, were conducted at weeks 18 and 19, respectively.

All animal procedures were done in compliance with the Animal Ethics/Research Committee of the University of Barcelona, and principles of laboratory animal care were followed (European and local government guidelines). All animals were randomly assigned to cohorts when used.

### *In vivo* assessment of glucose homeostasis

The intraperitoneal Glucose Tolerance Test (ipGTT) was performed by administering an intraperitoneal injection of D-glucose (2 g/Kg body weight). The fasting period prior to the test varied according to the dietary regimen: 5–6 hours for mice in physiological studies, and overnight for mice maintained on a high-fat diet. In both cases, glucose in tail vein blood was measured at the indicated time points using a clinical glucometer and Accu-Check test strips (Roche Diagnostics, Switzerland). Plasma samples were obtained by blood centrifugation and kept at -80°C for subsequent insulin determination by an Ultra-Sensitive Mouse Insulin ELISA Kit (Crystal Chem, IL, USA) (# Crystal Chem, Houston, TX, USA) following the manufacturer’s instructions.

The intraperitoneal Insulin Tolerance Test (ipITT) was performed after 4-5h of food deprivation by administration of an intraperitoneal injection of insulin at 0.75 IU/kg body weight (Regular Humulina® #710008.9, Lilly). Tail vein blood glucose levels were measured at 0, 15-, 30-, 60-, and 120-minute post-injection.

### Pancreatic immunofluorescence staining and morphometric analysis

Pancreata was harvested, fixed overnight in 10% (v/v) formalin, and subsequently washed, dehydrated and embedded in paraffin wax. For immunofluorescence in paraffin sections, a standard immunodetection protocol was followed as described elsewhere^17^. Briefly, tissues were rehydrated and, when required, subject to heat-mediated antigen retrieval in citrate buffer. After a blocking step in 5% donkey or goat serum, and permeabilization using 1% (v/v) Triton X-100, tissue sections were incubated overnight with insulin, glucagon and somatostatin antibodies and then for 1h with fluorescent-labeled secondary antibodies (Supplementary Table 2). Nuclei were stained with Hoechst 33258 (1/500, Sigma). Images were collected using a Leica AF6000 motorized system (Leica Microsystems, Manheim, Germany) equipped with a DMI6000 microscope, a high-resolution monochrome Hamamatsu Orca ER C4742-80 Digital Camera and a mercury metal halide bulb Leica EL6000 as light source. Image acquisition program was LASX Navigator and images obtained were subsequently analyzed using ImageJ/Fiji software (National Institutes of Health, Bethesda, MD, USA; https://imagej.net/ij/).

For histomorphometry analysis of endocrine cell mass, at least six non-consecutive (150µm apart) 4µm thick sections were analyzed per pancreas. Beta, alpha, and delta-cell fractional areas were quantified as insulin, glucagon and somatostatin-positive areas (respectively) relative to total pancreatic area. For each endocrine compartment, cell mass was calculated as the cell fractional area multiplied by pancreas weight.

### Intra-islet macrophage quantification

To analyze macrophage infiltration, pancreatic tissue sections were incubated overnight with insulin and F4/80 primary antibodies, and then for 1h with fluorescent-labeled secondary antibodies (Supplementary Table 2). Nuclei were stained with Hoechst 33258 (1/500, Sigma). Images were acquired using a Leica Thunder inverted widefield microscope (DMi8, Leica Microsystems, Manheim, Germany). Ilumination was provided by a pE-800 MB LED light source (CoolLED), and the DFT5 and CYR7 filter sets. Images were captured with a Leica K8 camera, using a 40x objective. Large-area images were obtained by automated stage scanning and stitched using the Smooth method in LAS X. Instant Computational Clearing (Thunder) was applied for background removal and deconvolution, using a feature size parameter of 8.51682µm and a strength of 0.9. Collected images were subsequently analyzed using ImageJ/Fiji software (National Institutes of Health, Bethesda, MD, USA; T http://rsb.info.nih.gov/ij/). To quantify islet inflammation, an average of 42 islets from 4 mice per genotype were analyzed, and the number of F4/80 positive cells inside each islet was manually counted. The percentage of islets containing 0, 1-3, 4-9, and equal or more than 10 cells, were represented.

### Islet studies

Mouse islets were obtained by collagenase digestion and Histopaque gradient purification as previously described^45^. Isolated islets were allowed for an overnight recovery in RPMI-1640 media with 11mM glucose, 10% fetal bovine serum, and antibiotics before performing experiments. To evaluate glucose-stimulated insulin secretion *in vitro*, islets were washed with phosphate-buffered saline (PBS) and preincubated for 30-45 min at 37°C in Hepes-buffered Krebs-Ringer buffer with 2mM glucose. Preincubation solution was then removed and islets subsequently incubated with the same buffer containing low (2mM) and high (20Mm) glucose concentrations for 60 min. Supernatants were collected after each incubation period to measure secreted insulin. Islet pellets were lysed in acid-alcohol-containing buffer (ethanol 75%, HCl 1.5%) and sonicated to determine insulin content. Insulin was measured using the Ultra-Sensitive Mouse Insulin ELISA Kit also used for plasma measurements. Fractional insulin release was calculated as percentage of insulin secreted relative to total content of the islets.

### Plasma and serum measurements

Blood was collected via the tail vein into EDTA-coated capillary tubes (NC9141704, Sarstedt) and subsequently centrifuged at 3,600 × rpm for 20 minutes at 4°C to obtain plasma. Serum samples were individually obtained at endpoint by centrifugation of the whole blood obtained by cardiac puncture at 8,000 × *g* for 15 min at 4°C twice. Both plasma and serum samples were stored at -80°C until further analysis.

Plasma free fatty acids (FFA) were measured using the ACS-ACOD enzymatic colorimetric method (WAKO; HR Series NEFA-HR(2)-R1, Cat# 434-91795 and HR Series NEFA-HR(2)-R2, Cat# 436-91995). Plasma triglycerides (TAG) were quantified using the GPO-POD enzymatic colorimetric assay (Spinreact Cat# 1001312).

Plasma Wisp1 levels were determined using a specific ELISA kit (Quantikine® ELISA #MWSP10, Biotechne-R&D Systems). Serum levels of leptin (#90030, Cristal Chem), Adiponectin (KE10044, Proteintech), IL-6 (Quantikine® ELISA #M6000B-1, R&D Systems), TNF-alpha (Quantikine® ELISA #MTA00B-1, R&D Systems) and CCL2/MCP-1 (KE10006, Proteintech) were measured using the respective commercial ELISA kits following the manufactureŕs instructions.

### RNA isolation and gene expression analysis

RNA from islets was isolated using NucleoSpin®RNA (Macherey-Nagel, Düren, Germany) following the manufacturer’s manual. Peripheral tissues were harvested, immediately snap-frozen in liquid nitrogen and stored at -80°C until processed. Total RNA was extracted from frozen tissue using TRI Reagent® (Sigma-Aldrich, Missouri, USA) according to the manufacturer’s protocol and then cleaned and DNAse-treated using DNase (1U/ul), (DNase I, Amplification Grade, Invitrogen) prior to cDNA synthesis. RNA concentration and purity were measured with a Nanodrop spectrophotometer (Thermo Fisher Scientific). First-strand cDNA was prepared using the Superscript III RT kit and random hexamer primers (Invitrogen, Carlsbad, CA, USA) in a total volume of 20 ul. Reverse transcription reaction was carried for 90 min at 50°C and an additional 10 min at 55°C and 15 min at 70°C.

Quantitative PCR was performed on an QuantStudio™ 5 detection system (Applied Biosystems, ThermoFisher Scientific) using GoTaq® qPCR Master Mix (Promega Biotech Ibérica, Alcobendas, Madrid, Spain). Gene expression was normalized to housekeeping genes (specified in the figure legends) and calculated using the ΔCt method, expressed as 2^(-ΔCt) unless otherwise indicated. Primer sequences are listed in Supplementary Table 1.

### Adipose tissue morphometry

Epididymal white adipose tissue (eWAT), and subcutaneous white adipose tissue (scWAT), were harvested and fixed overnight at 4°C with 10% neutral buffered formalin (HT501128-4L, Sigma-Aldrich). After fixation, tissues were washed in PBS, dehydrated through a graded series of ethanol solutions (70%, 80%, 95% and 100% for 1 hour each), cleared in xylene and embedded in paraffin wax using standard histological procedures. Serial sections were cut at a thickness of 2-3µm using a microtome and mounted on SuperFrost Plus microscope slides. Sections were then stained with Hematoxylin and Eosin (H&E) following standard protocols. Images were acquired using an upright microscope with Nikon’s CFI60 Infinity Optical Systemlight (Nikon Eclipse NiU).

For histomorphometry analysis, at least two non-consecutive (100µm apart) sections were analyzed per adipose depot and mouse. Cell size was quantified using an automated image process (in house-built macro called adiposize) in ImageJ/Fiji (National Institutes of Health, Bethesda, MD, USA; https://imagej.net/ij/) software.

### Macrophage infiltration in adipose tissue

Immunohistochemistry was performed using a Leica Bond RX platform for the anti-F4/80 D2S9R (Cell Signaling, 70076) at 1:1000 for 120 min. Prior to immunohistochemistry, paraffin-embedded sections were dewaxed and therefore epitope retrieval was performed with ER2 buffer (AR9640, Leica Biosystems). Washings were performed using the BOND Wash Solution 10x (AR9590, Leica). Quenching of endogenous peroxidase was performed by 10 min of incubation with Peroxidase-Blocking Solution at RT (S2023, Dako, Agilent). Non-specific unions were blocked using 5 % of goat normal serum (16210064, Life technology) mixed with 2.5 % BSA diluted in wash buffer for 60 min at RT. Secondary antibody used were the BrightVision Poly-HRP-Anti Rabbit IgG Biotin-free, ready to use (DPVR-110HRP, Immunologic). Antigen–antibody complexes were reveled with the DAB (Polymer) (Leica, RE7230¬CE). Sections were counterstained with hematoxylin (RE7107, Leica Biosystems) and mounted with Mounting Medium, Toluene-Free (CS705, Dako, Agilent) using a Dako CoverStainer. Specificity of staining was confirmed by staining with the rabbit IgG, polyclonal (Abcam, ab37415). Images were taken using an NDP scan 3.4.0 (C9600-12, Hamamatsu Photonics).

### Adipo-Clear and three-dimensional analysis of adipose tissue macrophages

Mice were anesthetized, and intracardiac perfusion was performed with 1xPBS followed by 4% PFA. Both eWAT and scWAT fat pads were harvested and post-fixed in 4% PFA at 4°C overnight, dehydrated with increasing concentrations of methanol in B1N buffer (20%, 40%, 60%, 80%, 30 min each) and delipidated with 100% dichloromethane (DCM) (three times, 30 min each). After delipidation, samples were rehydrated with decreasing concentrations of methanol in B1N buffer (100%, 80%, 60%, 40%, 20%, 30 min each), and subsequently permeabilized in PTwH buffer for 2 hours before antibody staining.

Samples were incubated for 6 days in primary antibody dilution (rat anti-mouse CD68, 1:400, BIO-RAD, MCA1957) in PTxwH buffer, followed by an incubation with a secondary antibody conjugated with Alexa-647 for 4 days (1:200, Jackson Immunoresearch, 712-605-153). Upon immunostaining, samples were embedded in agarose and subsequently dehydrated with increasing concentrations of methanol/H_2_O, washed in DCM, and cleared in Ethyl cinnamate for 48 hours (ECi, 112372 Sigma-Aldrich).

Whole-tissue samples were imaged using a light-sheet microscope (Ultramicroscope II, LaVision Biotec) equipped with a 1.3x objective lense. Samples were placed in an imaging reservoir filled with ECi and illuminated from the side by the 488 and 640nm lasers. Images were acquired with Inspector Pro software (LaVision BioTec). Three-dimensional video generation of the light sheet microscopy datasets were performed using the Napari Image Viewer (version 0.5.6) with the Napari-animation plugin. The images were down sampled at 50% of the original spatial resolution (5 um in all dimensions) to improve Napari’s performance, and only linear adjustments of brightness and contrast were applied for visualization purposes.

### Assessment of fibrosis in adipose tissue

Fibrosis was detected by using Masson’s trichrome stain (TRIC-KMA-10 Labkem), following manufacturer’s instructions. Tissue sections were observed under a Nikon Eclipse NI-U upright microscope (Niko-Europe B.V., Netherlands).

### Statistical information

Data were analyzed using GraphPad Prism 6.0 software and expressed as the mean ± standard error of the mean (SEM). The appropriate statistical test was determined based on the number of comparisons being made. Student’s *t*-tests (two tailed) were used for comparison of two groups. One-way ANOVA was used for analysis of three or more experimental groups. Two-way ANOVA was used for analysis of mouse serum Wisp1 concentrations and metabolic tests *in vivo* experiments. When finding significant results in ANOVA, recommended post-hoc tests (Tukey or Sidak) were applied to determine significant differences among multiple comparisons. p<0.05 was deemed significant. P values greater than 0.05 and lower than 0.1 were considered trends, and exact values are shown in Figures.

## ACKNOWLEDGEMENTS

We gratefully acknowledge Julia Palà Badules and Candice Fournier for their technical support in specific experiments conducted during this study. We also thank the Advanced Optical Microscopy Unit of the *Universitat de Barcelona* (*CCiTUB*) for their assistance in image acquisition using the Leica AF6000 motorized system.

This research was supported by the Spanish Ministerio de Ciencia, Innovación y Universidades (PID2022-139450OB-I00) and Instituto Carlos III (PI19/00896) to R.G.; and CIBERDEM (DEM22_PIM05) and SED (VIII Ayuda SED a Proyectos de Investigación Básica en Diabetes dirigidos por Jóvenes Investigadores) to R.F.R. P.C. was supported by NIDDK R01 DK120649. M.P. was recipient of a FPU fellowship (FPU22/01454) from the Spanish Ministerio de Ciencia, Innovación e Universidades. C.K.H. was supported by a Medical Scientist Training Program grant from the National Institute of General Medical Sciences of the National Institutes of Health under award number T32GM152349 to the Weill Cornell/Rockefeller/Sloan Kettering Tri-Institutional MD-PhD Program.

