## Supplemental data for "Uncovering the function of Wisp1 in whole-body glucose homeostasis: insights from Wisp1 knockout mice"

### for

**Includes:**

Figures S1-S10

Video captions: S1, S2

Tables S1, S2

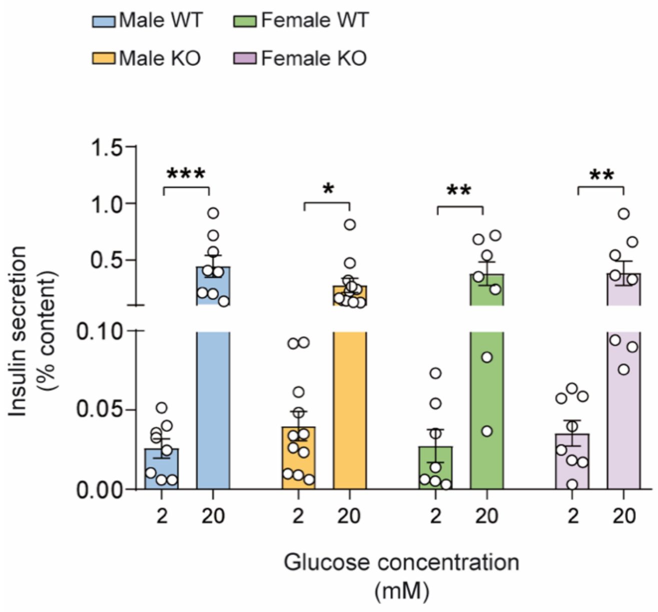


**Figure S1.** ***In vitro*** **insulin secretion by isolated islets from male and female Wisp1 WT and KO mice.**

Islets were isolated from 11-week-old male and female Wisp1 WT and KO mice. Results are depicted as fractional insulin release at low (2mM) and high (20mM) glucose concentration. Data are presented as mean±SEM from n=35-60 islet batches from 7-12 mice per sex and genotype. *$p<0.05$, **$p<0.01$, ***$p<0.001$. Comparisons were made using two-way ANOVA.

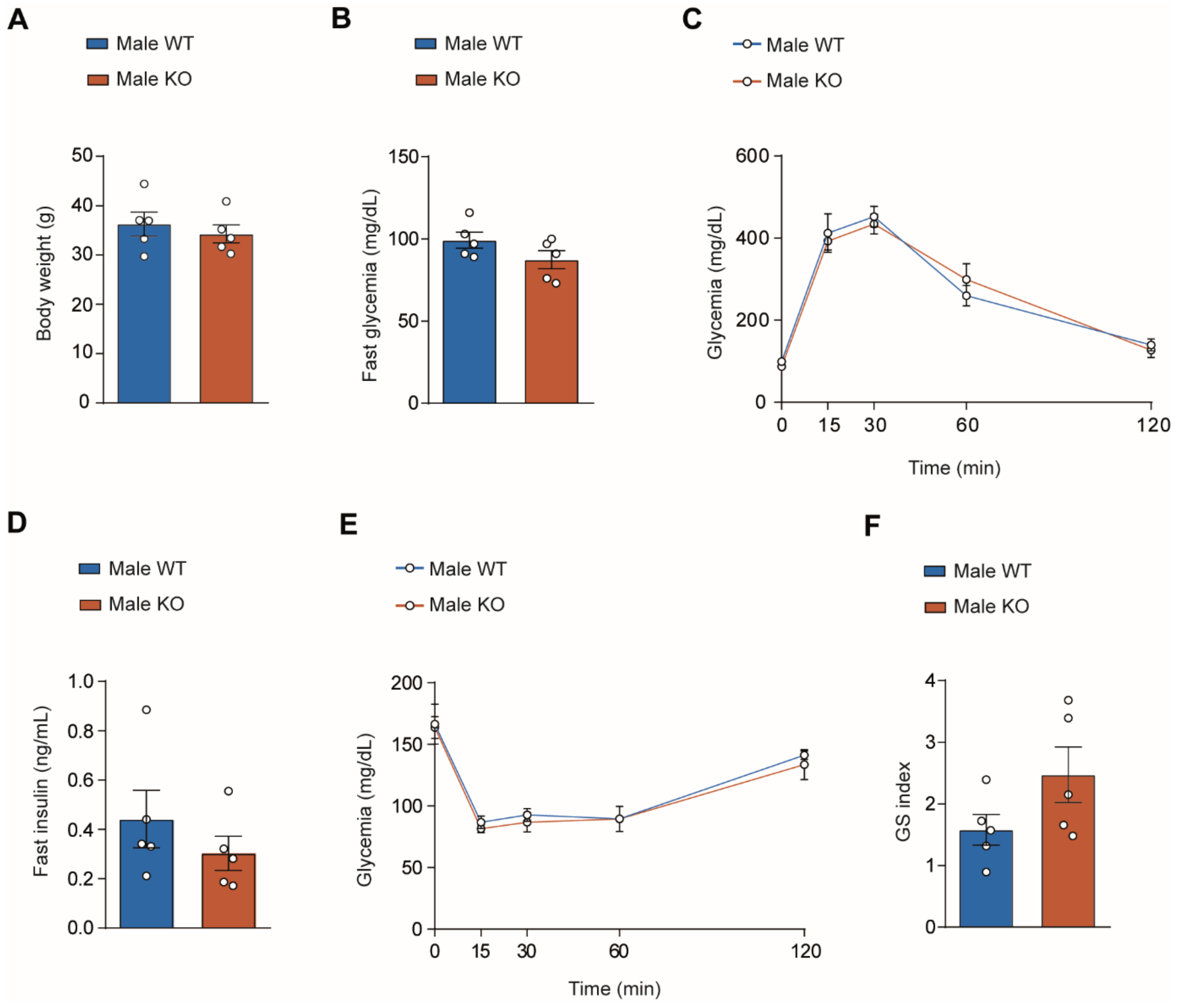


**Figure S2. Glucose homeostasis in 50-week-old male Wisp1 KO and WT mice.**

**A.** Body weight of 50-week-old male Wisp1 WT and KO mice. Data are presented as mean±SEM for n=5 mice, shown as individual points. Comparisons were made using two-tailed Student´s *t* test.

**B.** Overnight fasting glycemia of 50-week-old male Wisp1 WT and KO mice. Data are presented as mean±SEM for n=5 mice per genotype, shown as individual points. Comparisons were made using two-tailed Student´s *t* test.

**C.** ipGTT results in 50-week-old male Wisp1 WT and KO mice. Data are shown as mean±SEM for n=5 mice. Comparisons were made using a two-way ANOVA.

**D.** Overnight fasting plasma insulin levels of 50-week-old male Wisp1 WT and KO mice. Data are presented as mean±SEM for n=5 mice per genotype, shown as individual points. Comparisons were made using two-tailed Student´s *t* test.

**E.** ipITT results in male Wisp1 WT and KO mice at 50 weeks of age. Data are shown as mean±SEM for n=5 mice per genotype. Comparisons were made using a two-way ANOVA.

**F.** *In vivo* glucose-stimulated insulin index (GS index), calculated as insulin levels 15 minutes after glucose injection relative to basal insulin levels following an overnight fasting in 50-week-old male Wisp1 WT and KO mice. Data are presented as mean±SEM for n=5 mice, shown as individual points. Comparisons were made using two-tailed Student´s *t* test.

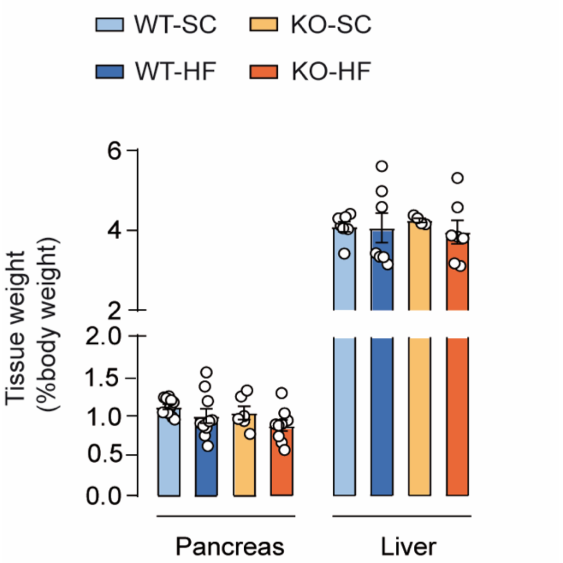


**Figure S3. Tissue weights in male Wisp1 KO mice under HF diet conditions.**

Pancreata and livers were harvested from male Wisp1 WT and KO mice fed either a HF or a SC diet for 20 weeks. Results are expressed as percentage of the tissue weight relative to total body weight. Data are shown as mean±SEM for=6-11 mice, shown as individual points. Comparisons were made using one-way ANOVA.

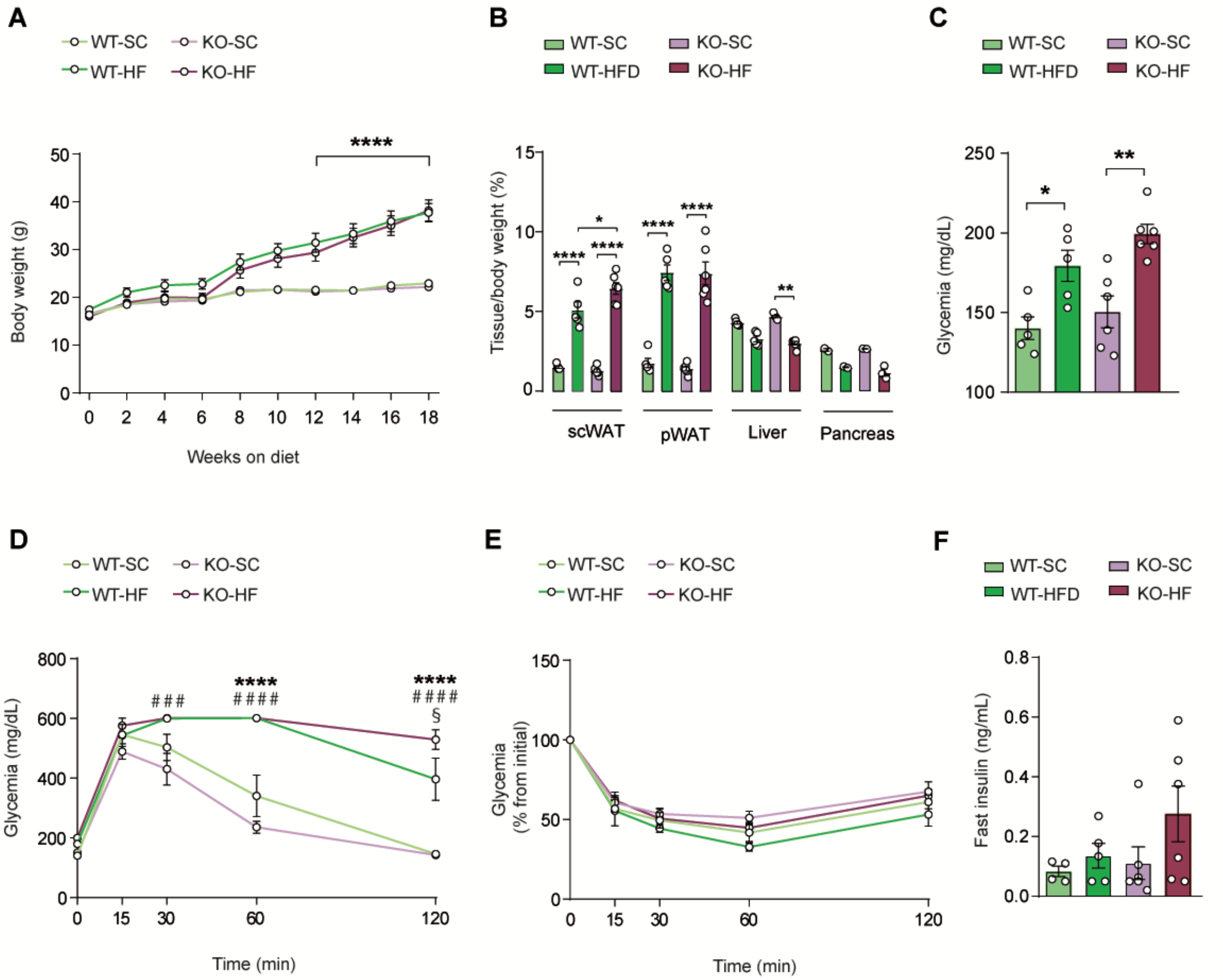


**Figure S4. Glucose homeostasis in female Wisp1 KO mice under HF diet conditions.**

- **A.** Body weight was monitored in female WT and KO mice fed either a HF or a SC diet for 20 weeks. Data are presented as mean±SEM from n=5-6 mice per group. ****p<0.0001 Comparisons were made using two-way ANOVA.
- **B.** Comparison of the relative weight of visceral (perigonadal, pWAT) and subcutaneous (scWAT) adipose tissue, as well as liver and pancreas, across the different diet and genotype groups. Data are shown as mean±SEM from n=2-6 mice, shown as individual points. *p<0.05, **p<0.01, ****p<0.0001. Comparisons were made using one-way ANOVA within each depot.
- **C.** Overnight fasting glycemia of in female WT and KO mice fed either a HF or a SC diet for 19 weeks. Data are presented as mean±SEM from n=5-6, shown as individual points. *p<0.05, **p<0.01. Comparisons were made using one-way ANOVA.
- **D.** ipGTT results in female Wisp1 WT and KO mice fed either a HF or a SC diet for 19 weeks. Data are shown as mean±SEM from n=5-6 per group. ###p<0.001 (KO-SC *vs* KO-HF), ****p<0.0001 (WT-SC *vs* WT-HF), #### p<0.0001 (KO-SC *vs* KO-HF), § p<0.05 (WT-HF *vs* KO-HF). Comparisons were made using two-way ANOVA.
- **E.** ipITT results in female Wisp1 WT and KO mice fed either a HF or a SC diet for 18 weeks. Results are expressed as the percentage of variation from initial glucose concentration (100%). Data are presented as the mean±SEM for n=5-6 mice per group. Comparisons were made using two-way ANOVA.
- **F.** Plasma insulin levels after an overnight fast in male Wisp1 WT and KO mice fed either a HF or a SC diet for 19 weeks. Data are presented as mean±SEM from n=4-6 mice, shown as individual points. Comparisons were made using one-way ANC[illegible]

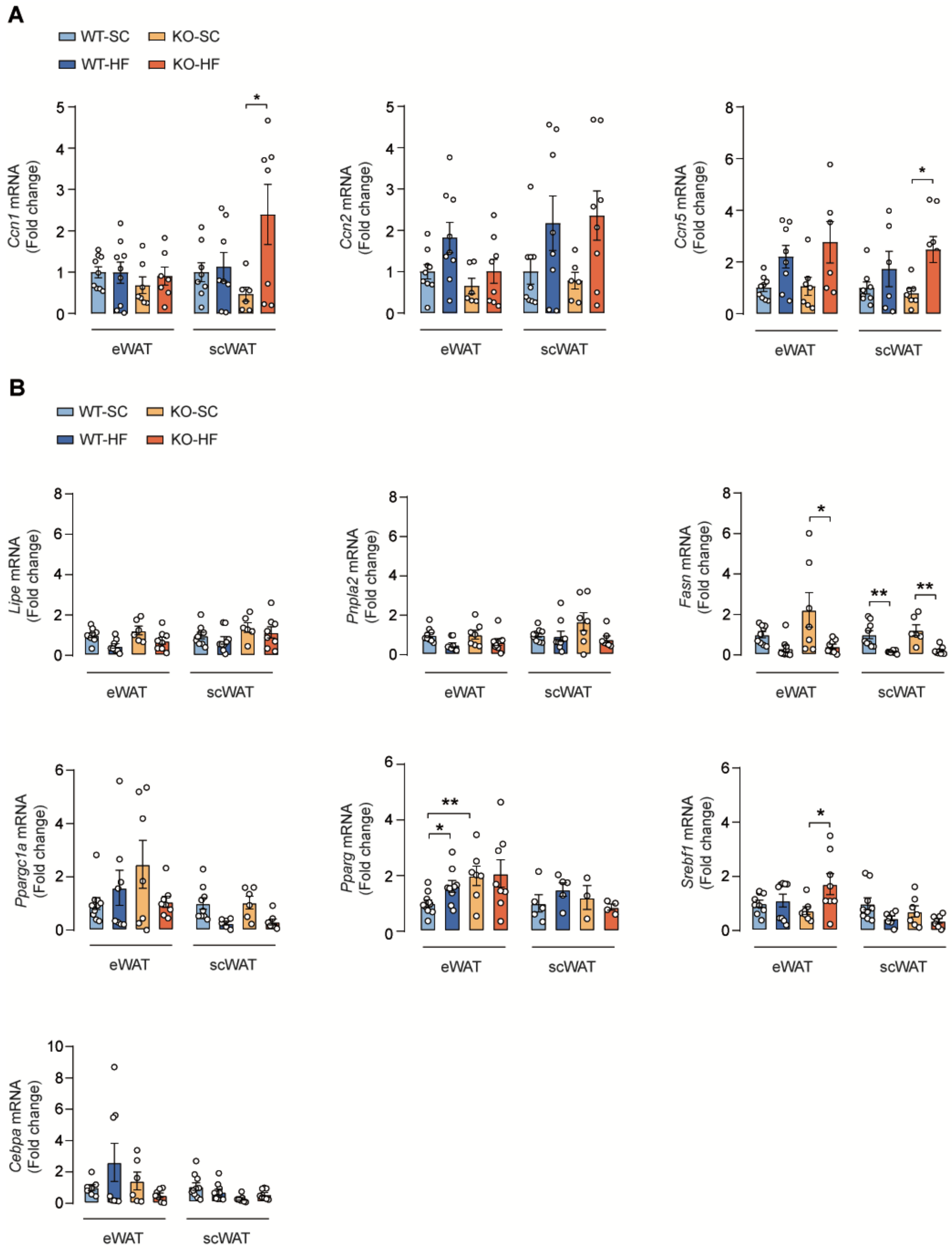


**Figure S5. Gene expression profile in scWAT and eWAT of Wisp1 KO mice under HF conditions.**

**A.** Gene expression of other members of the CCN family of proteins in the indicated adipose tissue depot in male Wisp1 WT and KO mice fed either a HF or a SC diet for 20 weeks. Expression was assessed by qPCR, using *Rplp0* and *Tbp* as housekeeping genes. Results are expressed relative to WT-SC, given the value of 1. Data are shown as mean±SEM from n=5-9 mice, shown as individual points. *p<0.05. Comparisons were made using a one-way ANOVA.

**B.** Gene expression of selected adipocyte differentiation, lipolisis and lipogenic markers in the indicated adipose tissue depot in male Wisp1 WT and KO mice fed either a HF or a SC diet for 20 weeks. Expression was assessed by qPCR, using *Rplp0* and *Tbp* as housekeeping genes. Results are expressed relative to WT-SC, given the value of 1. Data are presented as mean±SEM from n=4-9 mice, shown as individual points. *p<0.05, **p<0.01. Comparisons were made using a one-way ANOVA.

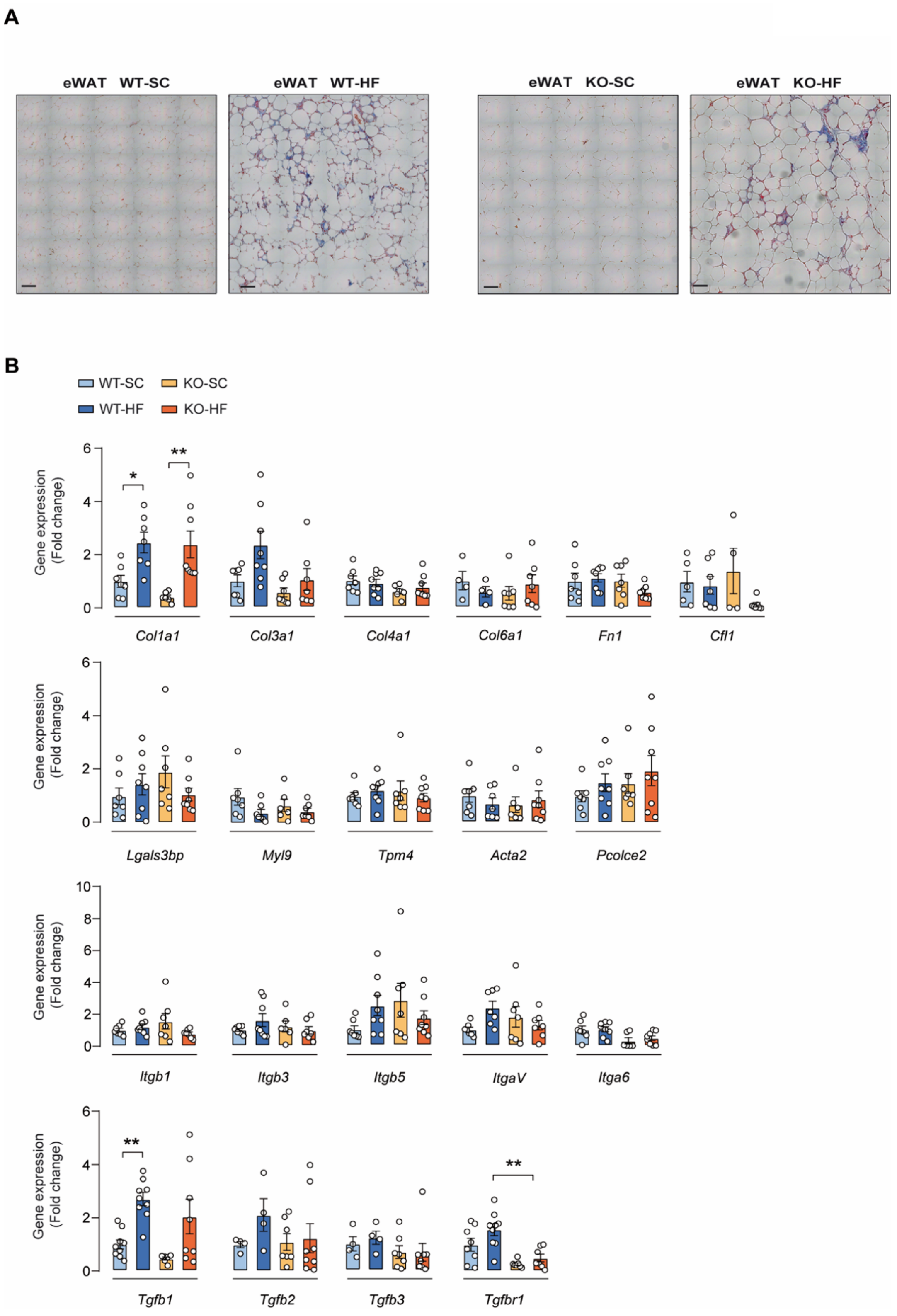


**Figure S6. Assessment of eWAT fibrosis in male Wisp1 KO mice under HF diet conditions.**

**A.** Representative Masson´s trichrome staining of eWAT from male Wisp1 WT and KO mice fed a HF diet for 20 weeks. Scale bar: 100µm.

**B.** Gene expression of fibrosis-related genes in male Wisp1 WT and KO mice fed either a HF or a SC diet for 20 weeks. Expression was assessed by qPCR, using *Rplp0* and *Tbp* as housekeeping genes. Results are expressed relative to WT-SC, given the value of 1. Data are shown as mean±SEM from n=4-9 mice, shown as individual points. *p<0.05, **p<0.01. Comparisons were made using a one-way ANOVA.

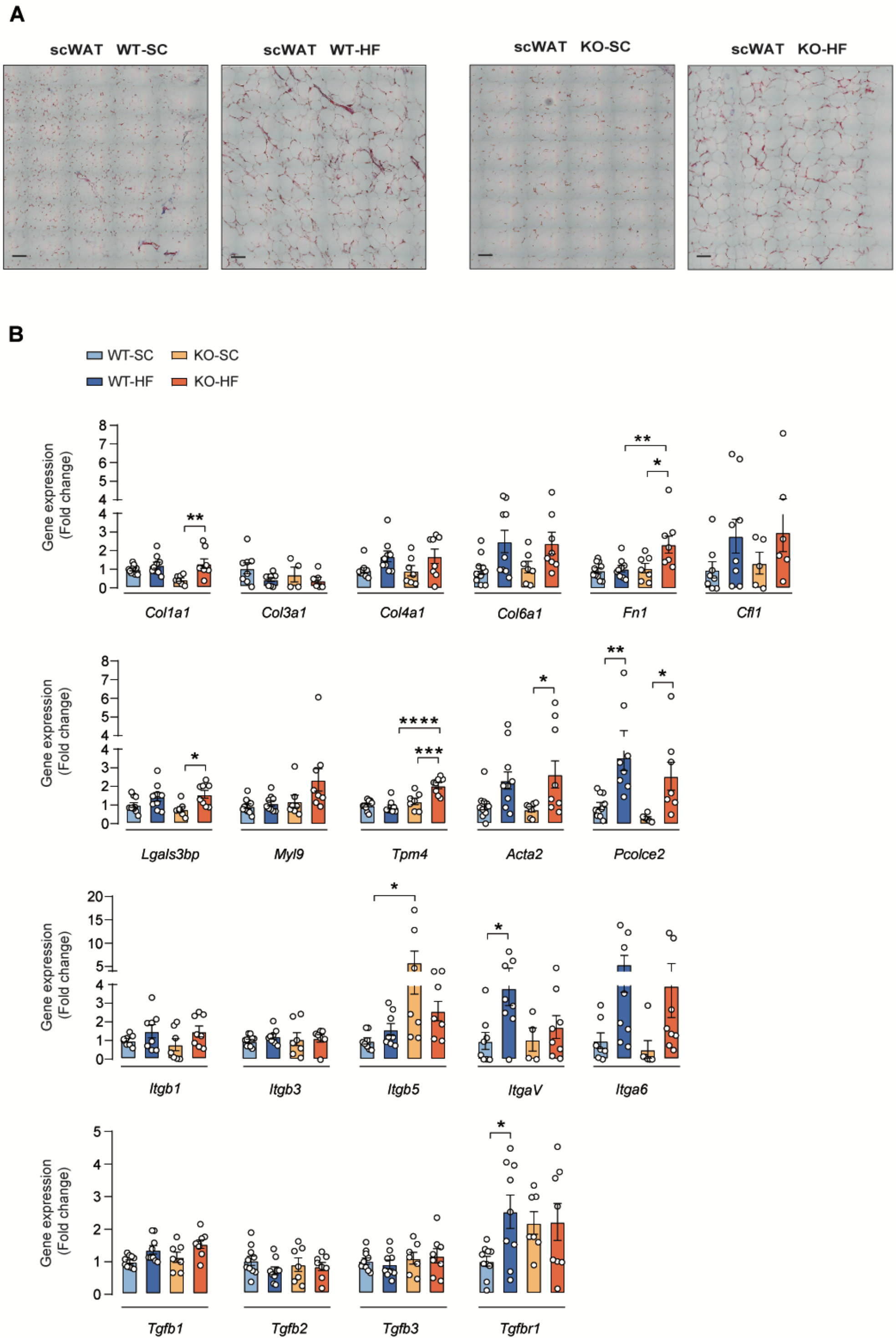


**Figure S7. Assessment of scWAT fibrosis in male Wisp1 KO mice under HF diet conditions.**

**A.** Representative Masson´s trichrome staining of scWAT from male Wisp1 WT and KO mice fed a HF diet for 20 weeks. Scale bar 100μm.ç

**B.** Gene expression of fibrosis-related genes in male Wisp1 WT and KO mice fed either a HF or a SC diet for 20 weeks. Expression was assessed by qPCR, using *Rplp0* and *Tbp* as housekeeping genes. Results are expressed relative to WT-SC, given the value of 1. Data are shown as mean±SEM from n=4-9 mice, shown as individual points. *p<0.05, **p<0.01, ***p<0.001, ****p<0.0001. Comparisons were made using a one- way ANOVA.

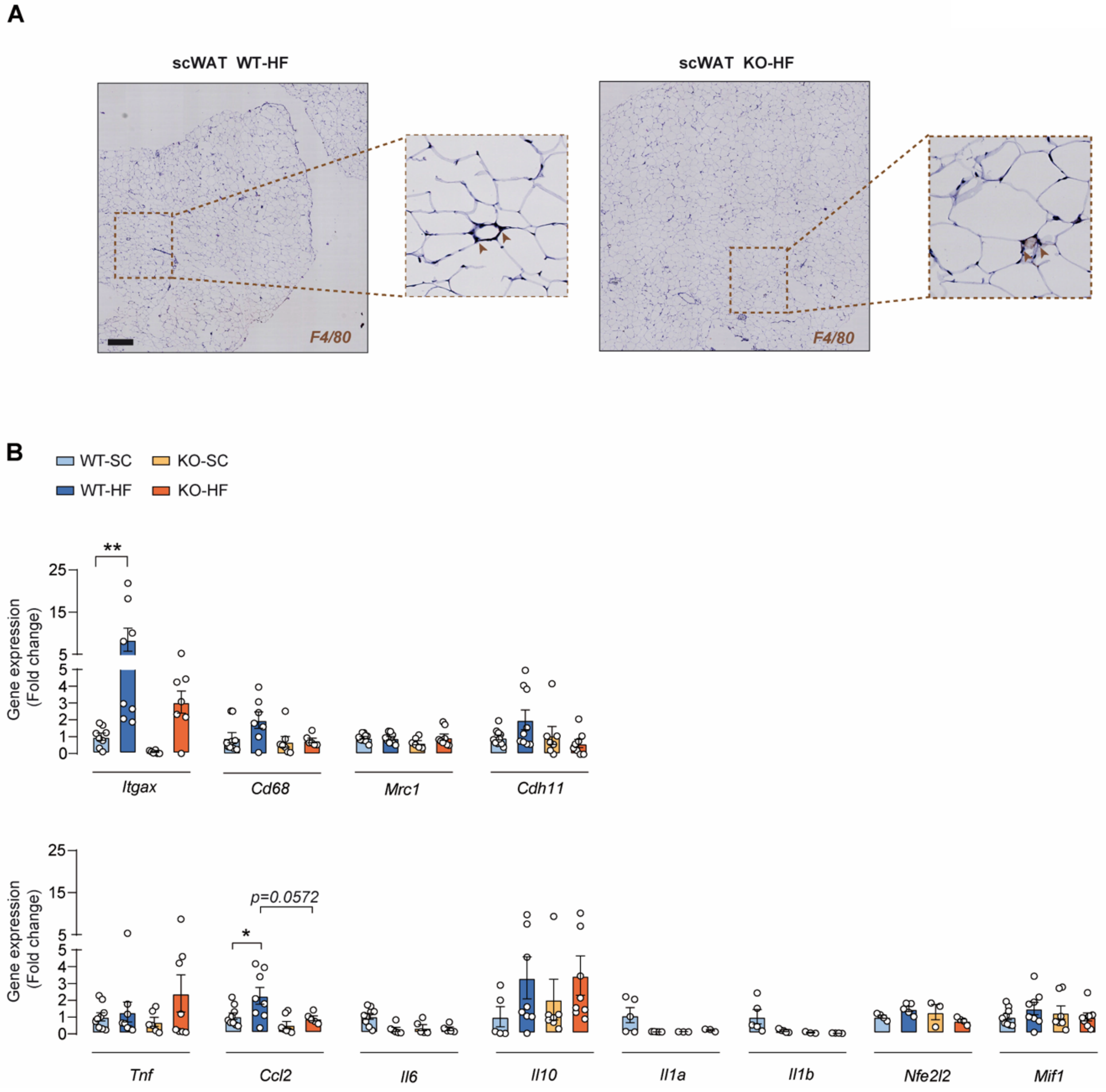


**Figure S8. Assessment of scWAT inflammation in male Wisp1 KO mice under HF diet conditions.**

**A.** Representative immunohistochemistry images showing positive F4/80 staining in scWAT from male Wisp1 WT and KO mice fed a HF diet for 20 weeks. Scale bar 500µm.

**B.** Gene expression of inflammation-related genes in male Wisp1 WT and KO mice fed either a HF or a SC diet for 20 weeks. Expression was assessed by qPCR, using *Rplp0* and *Tbp* as housekeeping genes. Results are expressed relative to WT-SC, given the value of 1. Data are shown as mean±SEM from n=5-9 mice, shown as individual points. *p<0.05, **p<0.01. Comparisons were made using a one-way ANOVA.

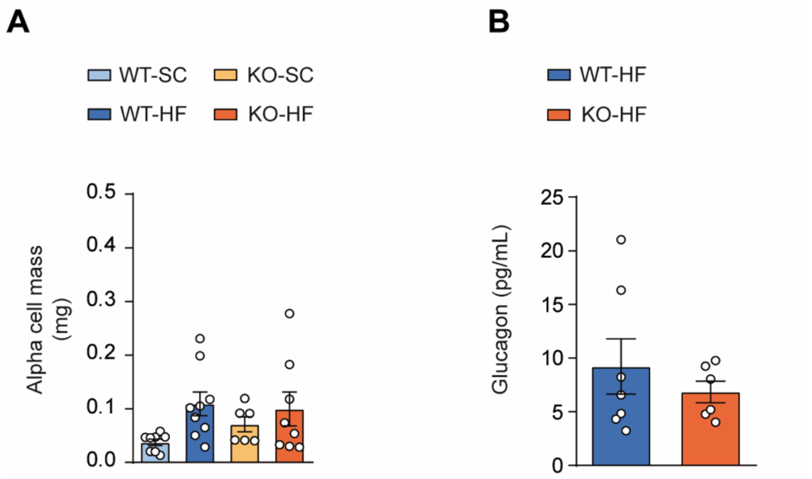


**Figure S9. Characterization of the alpha cell compartment in Wisp1 KO mice under HF conditions.**

**A.** Morphometric quantification of alpha-cell mass in fixed pancreatic sections from male Wisp1 WT and KO mice fed either a HF or a SC diet for 20 weeks. Data are shown as mean±SEM from n=6-9 mice, shown as individual points. **p<0.01. Comparisons were made using a one-way ANOVA.

**B.** Plasma glucagon levels after an overnight fast in male Wisp1 WT and KO mice fed either a HF or a SC diet for 20 weeks. Data are presented as mean±SEM for n=6-7 mice, shown as individual points. Comparisons were made using one-way ANOVA.

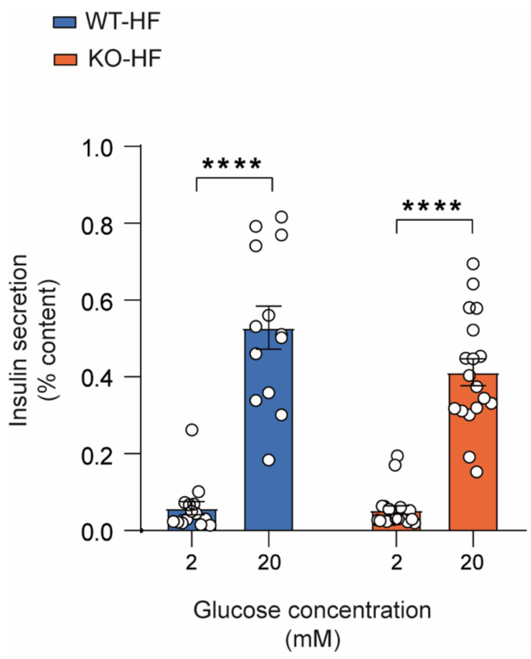


**Figure S10.** ***In vitro*** **insulin secretion by isolated islets from male Wisp1 WT and KO mice fed a high fat diet.**

Islets were isolated from male Wisp1 WT and KO mice fed a HF diet for 20 weeks. Results are depicted as fractional insulin release at low (2mM) and high (20mM) glucose concentration. Data are presented as mean±SEM from n=14-18 islet batches from 3 WT and 4 KO mice. *****$p<0.0001$. Comparisons were made using two-way ANOVA.

#### Video captions

**Video S1.** Representative two-dimensional reconstruction of eWAT tissue from a Wisp1 WT mouse, stained with the pan-macrophage marker Cd68. Reconstruction was made with Napari Image Viewer (version 0.5.6) with the Napari-animation plugin, down-sampling images by 50% for performance.

https://youtu.be/IN9sxuoJKKw

**Video S2.** Representative two-dimensional reconstruction of eWAT tissue from a Wisp1 KO mouse, stained with the pan-macrophage marker Cd68. Reconstruction was made with Napari Image Viewer (version 0.5.6) with the Napari-animation plugin, down-sampling images by 50% for performance.

https://youtu.be/aD6VMpKVeKs

**Table S1. List of oligonucleotides used for indicated techniques**

| Gene | Sequence (5′ to 3′) | Used in |
|---|---|---|
| *Wisp1Neo* | GTGCTTTACGGTATCGCCGCT | Genotyping |
| *Wisp1Exon2L* | ACCCCCACAACAATGACCT | Genotyping |
| *Wisp1Exon2R* | AGCTGCTGGGCACATATCTT | Genotyping |
| *mTbp-fw* | ACCCTTCACCAATGACTCCTATG | qRT-PCR |
| *mTbp-rv* | ATGATGACTGCAGCAAATCGC | qRT-PCR |
| *mRplp0-fw* | GAGGAATCAGATGAGGATATGGGA | qRT-PCR |
| *mRplp0-rv* | AAGCAGGCTGACTTGGTTGC | qRT-PCR |
| *mRpl32-fw* | GCTGCCATCTGTTTTACGG | qRT-PCR |
| *m Rpl32-rv* | TGACTGGTGCCTGATGAACT | qRT-PCR |
| *mCyr61-CCN1-fw* | AAAGGCAGCTCACTGAAG | qRT-PCR |
| *mCyr61-CCN1-rv* | GCCGGTATTTCTTGACAC | qRT-PCR |
| *mCtgf-CCN2-fw* | TTCCCGAGAAGGGTCAAGCT | qRT-PCR |
| *mCtgf-CCN2-rv* | TCCTTGGGCTCGTCACACA | qRT-PCR |
| *mWisp1-CCN4-fw* | GGTATCTCCACTCGGATCTCT | qRT-PCR |
| *mWisp1-CCN4-rv* | CCCTGCCTTGATGTGTAGTT | qRT-PCR |
| *mWisp2-CCN5-fw* | CCCCAGGAGAATACAGGTGC | qRT-PCR |
| *mWisp2-CCN5-rv* | GGCAGATGCAGGAGTGACAA | qRT-PCR |
| *mLep-fw* | ACATTTCACACACGCAGTCGG | qRT-PCR |
| *mLep-rv* | CAGCACATTTTGGGAAGGCAG | qRT-PCR |
| *mAdipoq-fw* | TGACGACACCAAAAGGGCTC | qRT-PCR |
| *mAdipoq-rv* | CACAAGTTCCCTTGGGTGGA | qRT-PCR |
| *mCfd-fw* | GTGCAAGTGAACGGCACACA | qRT-PCR |
| *mCfd-rv* | AGAGTCGTCATCCGTCACTCC | qRT-PCR |
| *mRtn-fw* | CTGTGGCTCGTGGGACATTC | qRT-PCR |
| *mRtn-rv* | CTCCCCGTCCCTGTCAACAT | qRT-PCR |
| *mItagx-fw* | GAGGCTGCAAGCATCATTCG | qRT-PCR |
| *mItagx-rv* | GCATCAAAGTTCTCCACGCTG | qRT-PCR |
| *mCd68-fw* | GTTACTCTCCTGCCATCCTTC | qRT-PCR |
| *mCd68-rv* | GCAGGGTTATGAGTGACAGTTG | qRT-PCR |
| *mMrc1-fw* | ACGAGCAGGTGCAGTTTACA | qRT-PCR |
| *mMrc1-rv* | ACATCCCATAAGCCACCTGC | qRT-PCR |
| *mCdh11-fw* | GAAATCTATCATGCCAATGTGC | qRT-PCR |
| *mCdh11-rv* | CACTATTTCCATAGGTGGGATCA | qRT-PCR |
| *mTnf-fw* | GTTGTACCTTGTCTACTCCCAG | qRT-PCR |
| *mTnf-rv* | GGTTGACTTTCTCCTGGTATGAG | qRT-PCR |

| | | |
|---|---|---|
| *mCcl2-fw* | CACTCACCTGCTGCTACTCA | qRT-PCR |
| *mCcl2-rv* | GCTTGGTGACAAAAACTACAGC | qRT-PCR |
| *mIl1a-fw* | CCCATGATCTGGAAGAGACCA | qRT-PCR |
| *mIl1a-rv* | CAAACTTCTGCCTGACGAGC | qRT-PCR |
| *mIl1b-fw* | TGCCACCTTTTGACAGTGATG | qRT-PCR |
| *mIl1b-rv* | TGATGTGCTGCTGCGAGATT | qRT-PCR |
| *mIl6-fw* | CAGAGGATACCACTCCCAAC | qRT-PCR |
| *mIl6-rv* | CAATCAGAATTGCCATTGCAC | qRT-PCR |
| *mIl10-fw* | AGGCGCTGTCATCGATTTCT | qRT-PCR |
| *mIl10-rv* | ATGGCCTTGTAGACACCTTGG | qRT-PCR |
| *mNfe2l2-fw* | CCGCTACACCGACTACGATT | qRT-PCR |
| *mNfe2l2-rv* | ACCTTCATCACCAACCCAAG | qRT-PCR |
| *mMif1-fw* | CAGAGGGGTTTCTGTCGGAG | qRT-PCR |
| *mMif1-rv* | GTGCACTGCGATGTACTGTG | qRT-PCR |
| *mLipe-fw* | GCGCTGGAGGAGTGTTTTT | qRT-PCR |
| *mLipe-rv* | CGCTCTCCAGTTGAACCAAG | qRT-PCR |
| *mPnpla2-fw* | TGACCATCTGCCTTCCAGA | qRT-PCR |
| *mPnpla2-rv* | TGTAGGTGGCGCAAGACA | qRT-PCR |
| *mFasn-fw* | CAAGTGTCCACCAACAAGCG | qRT-PCR |
| *mFasn-rv* | GGAGCGCAGGATAGACTCAC | qRT-PCR |
| *mPpargc1a-fw* | GGAGCCGTGACCACTGACA | qRT-PCR |
| *mPpargc1a-rv* | TGGTTTGCTGCATGGTTCTG | qRT-PCR |
| *mPparg-fw* | TGAAAGAAGCGGTGAACCACTG | qRT-PCR |
| *mPparg-rv* | TGGCATCTCTGTGTCAACCATG | qRT-PCR |
| *mSrebf1-fw* | ACTGGACACAGCGGTTTTGA | qRT-PCR |
| *mSrebf1-rv* | CTCAGGAGAGTTGGCACCTG | qRT-PCR |
| *mCebpa-fw* | TACCGAGTAGGGGGAGCAAA | qRT-PCR |
| *mCebpa-rv* | TCATTTTTCTCACGGGGCCA | qRT-PCR |
| *mCol1a1-fw* | GAACTGGACTGTCCCAACCC | qRT-PCR |
| *mCol1a1-rv* | CTTGGGTCCCTCGACTCCTA | qRT-PCR |
| *mCol3a1-fw* | CTGTAACATGGAAACTGGGGAAA | qRT-PCR |
| *mCol3a1-rv* | CCATAGCTGAACTGAAAACCACC | qRT-PCR |
| *mCol4a1-fw* | AGACTTTGCCCCAACAGGAG | qRT-PCR |
| *mCol4a1-rv* | TCTTTCCCAGGTTTTCCCCG | qRT-PCR |
| *mCol6a1-fw* | GCCGGTGATGAAGGAAATGC | qRT-PCR |
| *mCol6a1-rv* | GTCTCCCCTTGGTCCTCTCA | qRT-PCR |
| *mFn1-fw* | GATGTCCGAACAGCTATTTACCA | qRT-PCR |

| | | |
|---|---|---|
| *m Fn1-rv* | CCTTGCGACTTCAGCCACT | qRT-PCR |
| *mCfl1-fw* | CAGACAAGGACTGCCGCTAT | qRT-PCR |
| *mCfl1-rv* | TTGCTCTTGAGGGGTGCATT | qRT-PCR |
| *mLgals3bp-fw* | TGCTGGTTCCAGGGACTCAA | qRT-PCR |
| *mLgals3bp-rv* | CCACCGGCCTCTGTAGAAGA | qRT-PCR |
| *mMyl9-fw* | CTCTGCAGCAGGGAAACCC | qRT-PCR |
| *mMyl9-rv* | CTTCTTGGTGGTCTTGGCCT | qRT-PCR |
| *mTpm4-fw* | AAGCCGACCGCAAGTATGAG | qRT-PCR |
| *mTpm4-rv* | GTTCAGATACCTCCGCCCTC | qRT-PCR |
| *mActa2-fw* | CTACGAACTGCCTGACGGG | qRT-PCR |
| *mActa2-rv* | GCTGTTATAGGTGGTTTCGTGG | qRT-PCR |
| *mPcolce2-fw* | TGTGGCGGCATTCTTACCG | qRT-PCR |
| *mPcolce2-rv* | CCCTCAGGAACTGTGATTTTCCA | qRT-PCR |
| *mItgβ1-fw* | CAGGTGTCGTGTTTGTGAATG | qRT-PCR |
| *mItgβ1-rv* | GATCTGACCATTTGACGCTAGA | qRT-PCR |
| *mItgβ3-fw* | GGAATAGAACCCAGGACATCAC | qRT-PCR |
| *mItgβ3-rv* | CCGTATTTACTCTCGGCATCTT | qRT-PCR |
| *mItgβ5-fw* | GGATCAGCCAGAAGACCTTAAT | qRT-PCR |
| *mItgβ5-rv* | AATCTTCAGACCCTCACACTTC | qRT-PCR |
| *mItgaV-fw* | AGACATCCACTCCCTCTACAA | qRT-PCR |
| *mItgaV-rv* | AGTAGGTCATCTAGCCCATCTC | qRT-PCR |
| *mItga6-fw* | TGGACTCAGGGAAGGGTATT | qRT-PCR |
| *mItga6-rv* | CGCGGACTTCATGTCTCTTT | qRT-PCR |
| *mTgfβ1-fw* | GGTGGTATACTGAGACACCTTG | qRT-PCR |
| *mTgfβ1-rv* | CCCAAGGAAAGGTAGGTGATAG | qRT-PCR |
| *mTgfβ2-fw* | GGCTTTCATTTGGCTTGAGATG | qRT-PCR |
| *mTgfβ2-rv* | CTTCGGGTGAGACCACAAATAG | qRT-PCR |
| *mTgfβ3-fw* | CCACGAACCTAAGGGTTACTATG | qRT-PCR |
| *mTgfβ3-rv* | CTGGGTTCAGGGTGTTGTATAG | qRT-PCR |
| *mTgfβR1-fw* | TGTTTAATATTTGTCAGCATCCACCAG | qRT-PCR |
| *mTgfβR1-rv* | TGCACAAATACAATAAGACACAGAAAA C | qRT-PCR |
| *mTgfβR2-fw* | CAAGCCTCCAGAAGCCGTCCTCT | qRT-PCR |
| *mTgfβR2-rv* | GCTGCTGCTGCCGCTTCTAAACA | qRT-PCR |

**Table S2. List of antibodies**

IF:immunofluorescence
IHC: immunohistochemistry

| **Primary antibody** | **Raised in** | **Dilution** | **Source** |
|---|---|---|---|
| Insulin | guinea pig | 1/500 (IF) | DAKO #A0564 |
| Glucagon (K79bB10) | mouse | 1/1000 (IF) | Sigma #G2654 |
| Somatostatin | rabbit | 1/250 (IF) | DAKO #A0566 |
| F4/80 D2S9R | rabbit | 1/1000 (IHC)<br>1/50 (IF) | Cell Signaling #70076 |
| Rabbit IgG, polyclonal - Isotype Control | rabbit | 1/500 (IHC) | Abcam #ab37415 |
| Cd68 | rat | 1/400 (Adipoclear) | Biorad #MCA1957 |

| **Secondary antibody** | **Raised in** | **Dilution** | **Source** |
|---|---|---|---|
| Alexa Fluor®568 anti-guinea pig | goat | 1/400 (IF) | Thermo Scientific # A11075 |
| Alexa Fluor®488 anti-mouse | goat | 1/250 (IF) | Jackson Immunoresearch #115-545-166 |
| Alexa Fluor®488 anti-rabbit | donkey | 1/250 (IF) | Jackson Immunoresearch #711-546-152 |
| Alexa Fluor®647 anti-rabbit | donkey | 1/250 (IF) | Jackson Immunoresearch #711-605-152 |
| Alexa Fluor®647 anti-mouse | donkey | 1/250 (IF) | Jackson Immunoresearch #715-606-151 |
| Alexa Fluor®647 anti-rat | donkey | 1/200 (Adipoclear) | Jackson Immunoresearch #712-605-153 |
| Normal Donkey serum | | 5% (IF) | Jackson Immunoresearch #017-000-121 |
| Normal Goat Serum | | 5% (IF) | Jackson Immunoresearch #005-000-121 |
